# Force tuning explains changes in phasic dopamine signaling during stimulus-reward learning

**DOI:** 10.1101/2023.04.23.537994

**Authors:** Konstantin Bakhurin, Ryan N. Hughes, Qiaochu Jiang, Meghdoot Hossain, Boris Gutkin, Isabella P. Fallon, Henry H. Yin

## Abstract

According to a popular hypothesis, phasic dopamine (DA) activity encodes a reward prediction error (RPE) necessary for reinforcement learning. However, recent work showed that DA neurons are necessary for performance rather than learning. One limitation of previous work on phasic DA signaling and RPE is the limited behavioral measures. Here, we measured subtle force exertion while recording and manipulating DA activity in the ventral tegmental area (VTA) during stimulus-reward learning. We found two major populations of DA neurons that increased firing before forward and backward force exertion. Force tuning is the same regardless of learning, reward predictability, or outcome valence. Changes in the pattern of force exertion can explain results traditionally used to support the RPE hypothesis, such as modulation by reward magnitude, probability, and unpredicted reward delivery or omission. Thus VTA DA neurons are not used to signal RPE but to regulate force exertion during motivated behavior.

## Introduction

The ventral tegmental area (VTA) is the source of the mesolimbic dopamine (DA) pathway that has been implicated in reward and motivation ^1^. According to the influential reward prediction error (RPE) hypothesis, VTA DA neurons encode the difference between actual and predicted reward, providing a teaching signal for associative learning ^2,3^. However, others propose that DA contributes to vigor, incentive salience, or movement kinematics ^4–7^.

One reason for conflicting opinions on DA function is the lack of precise and continuous behavioral measurements. Many previous studies used head-fixed animals and behavioral measures that are usually limited to arm movements^8–10^ or licking ^3,11^. However, head restraint does not prevent mice from moving or attempting to move their head. In studies using freely moving animals, behavioral measures are limited to licking or time spent in the reward port ^12–16^. When continuous behavioral measures were used with high temporal and spatial resolution, DA activity in the substantia nigra pars compacta was found to be highly correlated with vector components of head velocity, regardless of reward prediction or outcome valence, and selective stimulation of DA neurons generated movements ^6^. Recent work using sensitive measures of force also showed that VTA DA neurons represent and regulate force exertion in head-fixed mice ^17,18^. These results suggest that previous observations in support of the RPE hypothesis could be explained by the contributions of DA signaling to online behavioral performance rather than learning.

In this study, we used force sensors to measure subtle movements in head-fixed mice during a Pavlovian stimulus-reward learning task previously used to study RPE signaling. Using optogenetics and *in vivo* electrophysiology, we showed that phasic DA activity in the VTA does not encode RPE, but is critical for modulating performance, in particular the force exerted by the animal.

## Results

Similar to previous work on RPE, mice were trained in a Pavlovian conditioning task: a conditioned stimulus (CS: 200 ms tone) is followed 1 second later by an unconditioned stimulus (US: sucrose reward) received just in front of the mouse (Figure 1A) ^3^. The head-fixation device has load cells that continuously measure forces exerted through the head-bar (Figure 1B)^17–20^. Such force exertion was not required to earn reward or to reach the spout, but reflect approach and consummatory behaviors. The reward spout was comfortably positioned ∼1 mm away from the lower jaw and was easily accessible with licking alone (**Supplemental Figure 1**).

**Figure 1.**
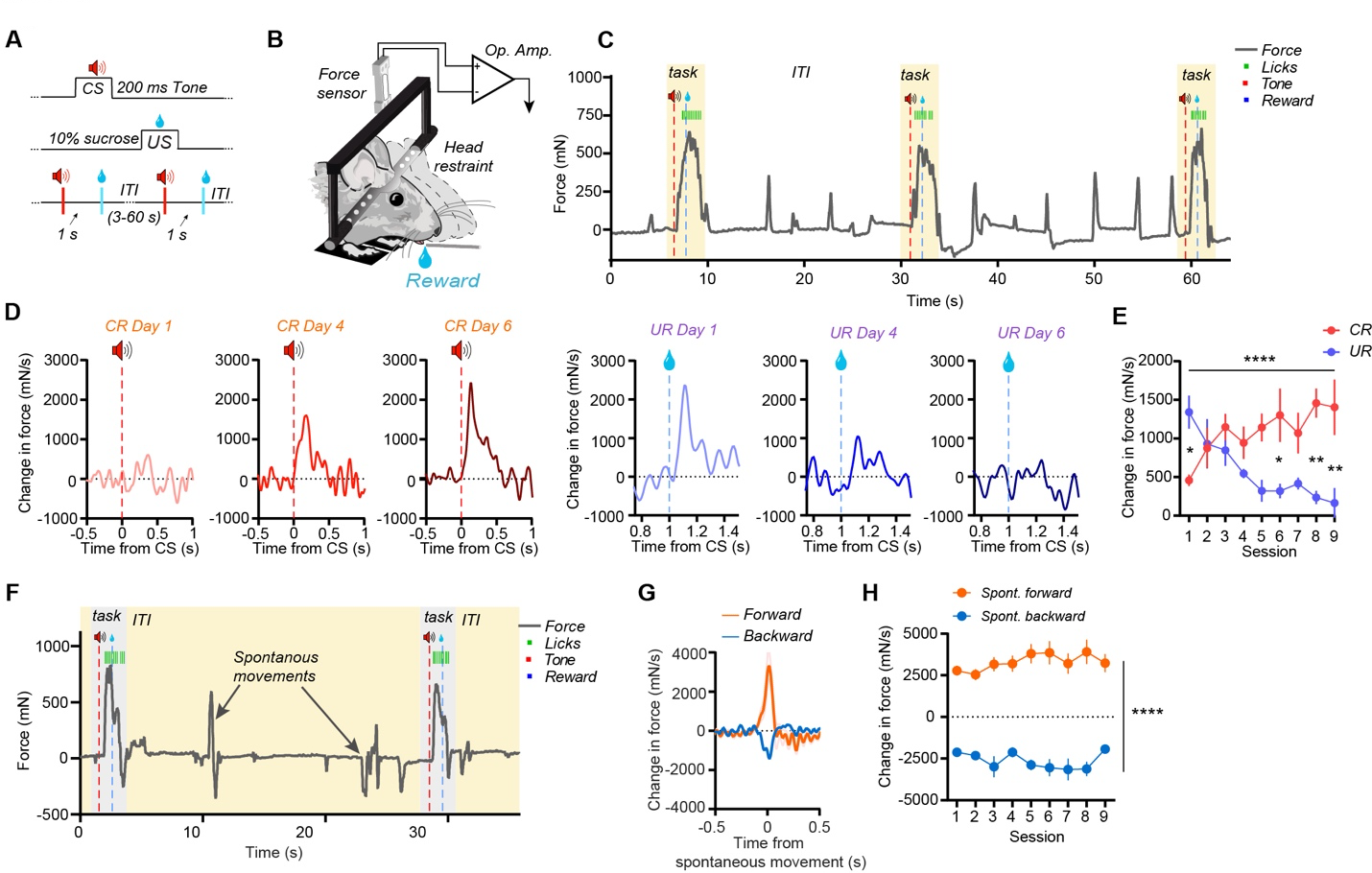
Force is a previously neglected behavioral measure in Pavlovian conditioning. **A)** Behavioral task design for stimulus-reward learning. **B)** Mice were restrained in a head-fixation device that continuously measures force exerted in the forward and backward directions. **C)** Representative force data obtained from a mouse well trained on a Pavlovian conditioning task. Note changes in force following CS and US presentations (**Supplemental Video 1**). **D)** Changes in force exertion following CS and US presentation across training in a representative mouse. **E**) Mice show a shift of peak force exertion from the UR to the CR during learning (Two-way ANOVA, significant interaction, F(8,28) = 8.0; Post-hoc tests found significant differences in force on days 1 (p<0.05), 6 (p<0.05), 8 (p<0.01) & 9 (p<0.01); n = 5 mice). **F)** Example force signal recorded between trials in a well-trained mouse. Note the presence of spontaneous backward and forward force exertion events during the ITI. **G**) Average spontaneous forward and backward change in force. **H**) No changes occurred in spontaneous movement amplitude over training (Two-way ANOVA, significant main effect of movement direction, F(1,18) = 171.4, significant interaction between training and direction on spontaneous force, F(8,134) = 2.65. Post-hoc tests showed that each session had significant differences between forward and backward movements (p < 0.0001), indicating the interaction reflected slight variability in differences in movement magnitude across sessions). Data points represent mean +/− SEM. * p<0.05, ** p < 0.01, **** p < 0.0001.

We could quantify changes in force exertion during stimulus-reward learning (**Supplemental Figure 1**). The mice exerted forward forces in anticipation of reward and applied additional force following reward delivery while they consumed the water (Figure 1C, **Supplemental video 1**). The patterns of force exertion changed systematically during training (Figure 1D). Force changes in response to the CS increased with training (Figure 1D, 1E), but those in response to the US decreased (Figure 1D, 1E). Thus, force exertion became more anticipatory during learning. Learning the task specifically involved changes in how force was exerted in response to CS and US. These subtle changes in movements have never been quantified in previous studies. They are not captured by standard measures of the conditioned response (CR) or unconditioned response (UR) such as licking (**Supplemental Figure 1**).

Mice also generated spontaneous movements during the inter-trial-interval (Figure 1F & G). While spontaneous movements became less frequent during training, their amplitudes did not change significantly, even in well-trained mice (Figure 1H, **Supplemental Figure 1**).

### Force tuning in VTA DA neurons

We recorded single unit activity from the VTA in mice with moveable optrodes and used optogenetic stimulation to confirm cell type (*n* = 1285 single units; n = 630 from putative DA neurons; *n* = 159 from tagged DA neurons; Figure 2A & **Supplemental Figures 2 & 3**)^17,18,20^. Because single units recorded during each session were counted independently, some of the single units may reflect repeated recordings from the same neurons.

**Figure 2.**
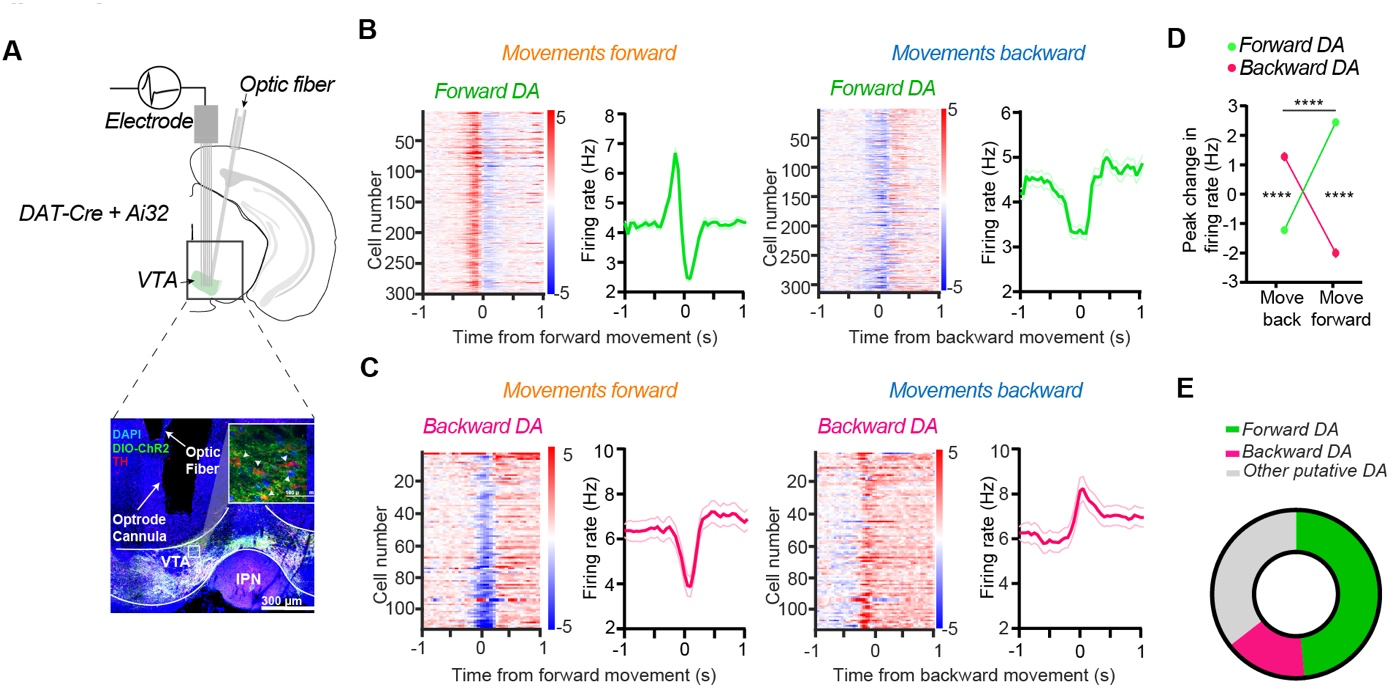
Opponent signaling of force direction by distinct DA neuron populations. **A)** *Top*, Identification of VTA dopamine (DA) neurons by optically tagging neurons that express *ChR2* only in DAT+ neurons. *Bottom*, co-registration of GFP with TH-expressing neurons. **B)** *Left*, Forward DA neurons are activated by spontaneous forward movements (n = 300 neurons). *Right*, the same Forward DA neurons show suppression in activity during spontaneous backward movement. C) *Left,* Backward DA neurons are suppressed by spontaneous forward movements (n = 116 neurons). *Right,* the same Backward DA neurons are activated by spontaneous movements backward. **D**) There was a significant interaction between the change in firing rate and the DA neuron class (Two-way ANOVA, F(1,408) = 930.13, p< 0.0001). Post-hoc tests found that peak firing of Forward DA neurons was greater than Backward DA during movements forward (p < 0.0001) and that peak firing of Backward DA was greater than Forward DA during movements backward (p < 0.0001). **E**) Forward and backward DA populations of the VTA make up approximately 65% of all DA neurons. Data represent mean +/− SEM. **** p < 0.0001.

We identified two major classes of DA neurons based on their unique direction-specific relationships to spontaneous force exertions (**Supplemental Figure 3**). One class of DA neurons (Forward DA, n = 300) produced a burst just prior to spontaneous forward movements (Figure 2B, **left**). In contrast, during spontaneous backward movements Forward DA neurons decreased their firing (Figure 2B, **right**). A second class of DA neurons (Backward DA, n = 116) showed the opposite pattern of activity, suppressing their firing pattern during spontaneous forward movements (Figure 2C, **left)** and increasing their firing (Figure 2C, **right**) during spontaneous backward movements. These two populations show opponent signaling depending on the direction of force exertion (Figure 2D). They made up ∼65% of confirmed DA neurons (Figure 2E). Many of the remaining neurons recorded in the VTA also showed activation to spontaneous movements (**Supplemental Figure 3**). Some had high firing rates and short-duration spike waveforms, properties associated with GABAergic projection neurons. Others were either not tagged and failed to show direction-specific force tuning. These neurons were not analyzed in this study.

During spontaneous movements, Forward and Backward neurons showed significant correlations with change in force in their preferred direction (Figure 3A & B). Representations of force by these populations were “rectified,” in that their firing rates had a linear relationship with force in their preferred direction but were uncorrelated with force in the opposite direction (Figure 3C & D).

**Figure 3.**
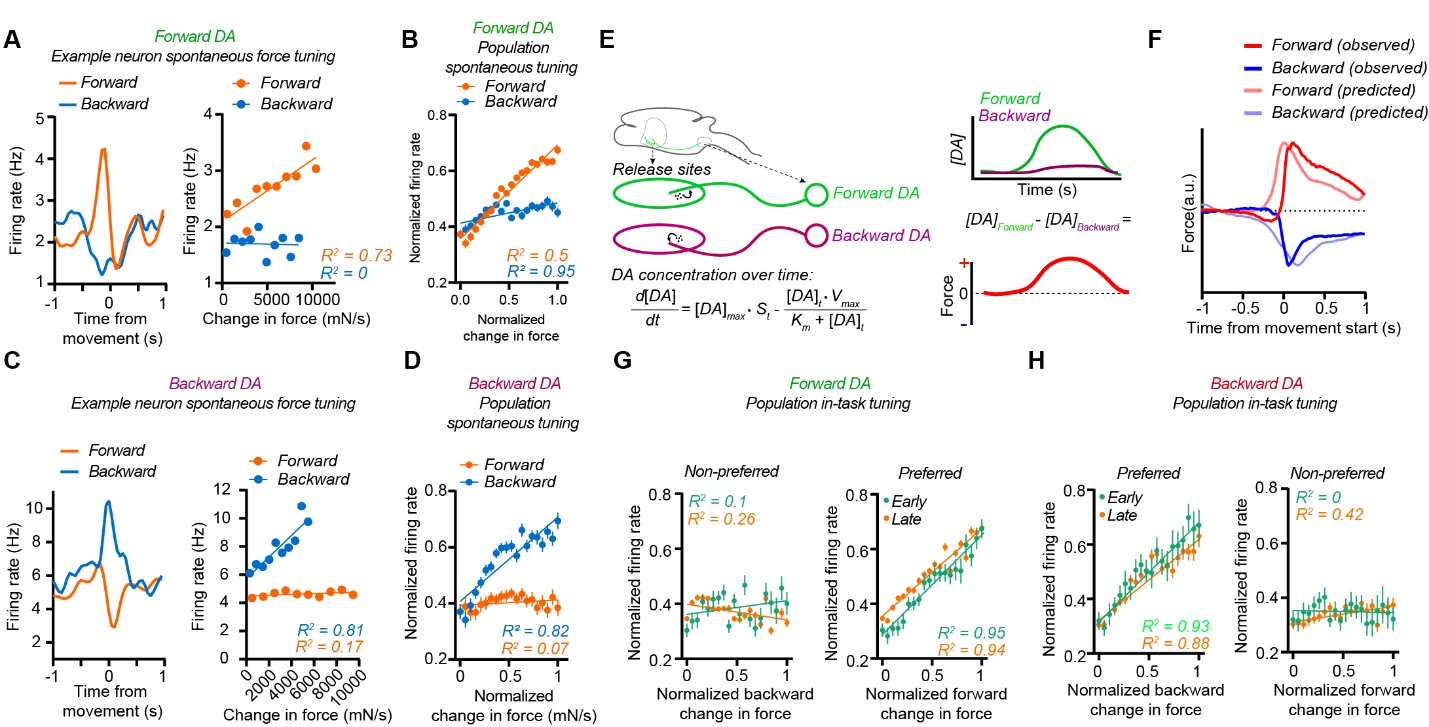
Force tuning by dopamine neurons is independent of learning. **A**) *Left*, representative Forward DA neuron showing preferred activity for spontaneous movements forward. *Right*, tuning profiles for the same Forward DA neuron, indicating a linear tuning for only forward movements (R^2^ for forward component = 0.73, p < 0.01, R^2^ for backward component: 0, p > 0.05). **B**) Forward DA neurons show a linear relationship with change in force magnitude during spontaneous forward movements but less so to spontaneous backward movements (R^2^ for forward = 0.94, p < 0.0001, R^2^ for backward = 0.5, p < 0.001, n = 300). **C)** *Left*, representative Backward DA neuron showing preferred activity for spontaneous movements backward. *Right*, tuning profile for the same Backward DA neuron (R^2^ for forward component = 0.17, p > 0.05, R^2^ for backward component: 0.81, p < 0.001). **D**) Backward DA neurons show linear relationship with change in force magnitude during spontaneous backward movements but not to spontaneous forward movements (R^2^ for forward = 0.07, p > 0.05, R^2^ for backward = 0.82, p < 0.0001, n = 116). **E**) Cartoon illustration of the modeling of DA concentration over time in the ventral striatum and its influence over force exertion. Extrasynaptic concentrations over time (d[DA]/dt) are determined by spike rate (S_t_) and reuptake kinetics (K_m_ & V_max_). Differences in concentration in target regions determine direction of force exertion. **F**) The average modeled DA concentration resulting from observed Forward and Backward DA population firing strongly resembles the force exerted by mice around movements (n = 156 Forward DA, 55 Backward DA, 4455 forward movements, 5313 backward movements). **G**) Forward DA neurons are preferentially tuned for forward movements during the task, regardless of learning (Preferred direction: R^2^ for Early = 0.95, p < 0.0001, n = 46; R^2^ for late = 0.94, p < 0.0001, n = 257. Non-preferred direction: R^2^ for Early = 0.1, p > 0.05, n = 46; R^2^ for Late = 0.26, p < 0.05, n = 257). **H**) Backward DA neurons are tuned for backward movements during the task regardless of learning (Preferred direction: R^2^ for Early = 0.93, p < 0.0001, n = 36; R^2^ for late = 0.88, p < 0.0001, n = 80. Non-preferred direction: R^2^ for Early = 0, p > 0.05, n = 36; R^2^ for Late = 0.42, p < 0.01, n = 80). Data represents mean +/− SEM.

If the activity of VTA DA neurons plays a causal role in movement, it must do so through DA receptor activation. Consequently, the dynamics of extracellular DA concentration should be related to the force measure. To model this relationship, we used standard DA release and reuptake parameter values ^21–24^ in a physiologically realistic biophysical model (Figure 3E). This model can predict DA concentration using the firing rates of DA neurons, using *[DA]_max_* (the instantaneous increase in DA concentration when DA neurons fire), *V_max_* (the maximal reuptake rate of DA in the striatum), and *K_m_* (the reciprocal of the affinity of the DA transporter). Using this model, we predicted the change in DA concentration in the target region (e.g. ventral striatum) over time resulting from spiking activity of Forward and Backward DA neurons. We assumed that these populations have different striatal regions that are involved in generating forces in opposite directions (Figure 3E). Because DA concentration is proportional to exerted force, the direction of force exerted is determined by the difference in DA concentrations in target regions. Within any tuning to fit the data or using any filters, the model predicted the direction of exerted force and its time course (Figure 3F). By varying the properties of the parameters *K_m_*, *V_max_*, and *[DA]_max_* within known biophysical constraints, it is possible to change the force profile (**Supplemental Figure 4**).

### Force tuning explains changes in phasic DA activity during learning

To determine whether these DA neurons show similar force tuning during the stimulus-reward task, we next analyzed force after CS and US presentation. Forward DA neurons again showed low correlations with backward movements that mice produced both early and late in learning (Figure 3G, Early: R^2^ = 0; Late: R^2^ = 0.26) but showed correlation with change in forward force (Early: R^2^ = 0.95; Late: R^2^ = 0.94). Backward DA neurons showed linear correlations with Backward force observed during the task (Figure 3H) but were less correlated with forward force. Together these results show that force tuning in VTA DA neurons was consistent across task-related and spontaneous movements.

DA neurons are known to show burst firing following CS presentation ^25,26^ during stimulus-reward learning and reduce their activity when rewards become better predicted. Such changes have been attributed to signaling an RPE ^3,8,26^. But our results suggest that changes in phasic DA may be explained by changes in performance rather than learning. In agreement with previous reports, Forward DA activity steadily increased during learning in response to the CS and decreased slightly after the US (Figure 4A & B). We found evidence in support of a shift of phasic DA activity from the US to the CS (Figure 4C). But these changes are accompanied by changes in the way force is exerted (Figure 1, Figure 4D & 4E). Initially there was more force exertion after reward delivery, but with training this behavior becomes more anticipatory, as the CR magnitude increased while the UR magnitude decreased (Figure 4F). These changes can explain the changes in the phasic activity of DA neurons during stimulus-reward learning.

**Figure 4.**
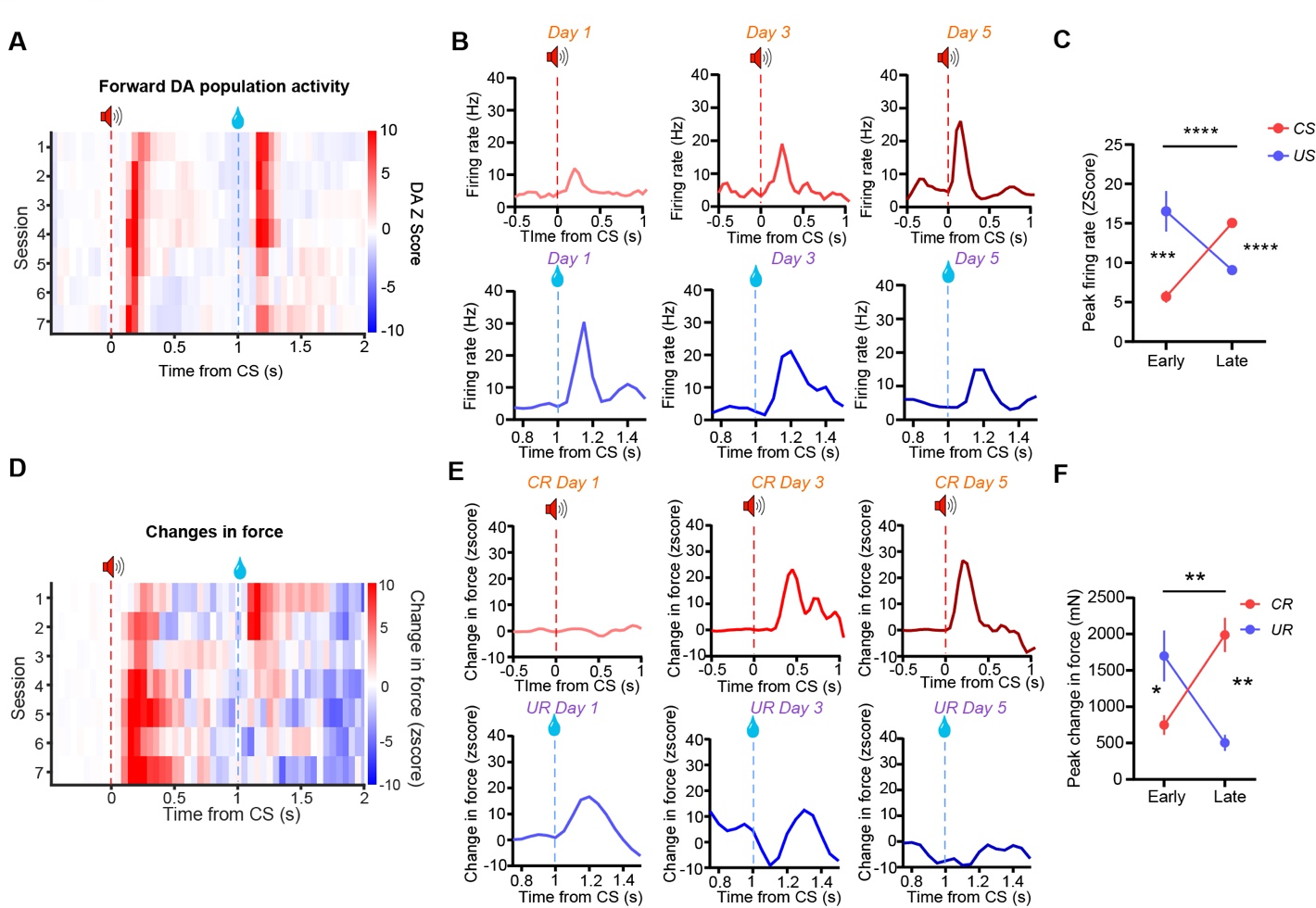
Force exertion explains shift in the timing of phasic DA activity from the US to the CS during learning. **A**) Population of Forward DA neurons recorded during learning. **B**) Forward DA population activity increases in response to the CS (*top row*) and decreases to the US (*bottom row*). **C**) Forward DA neurons show increases in rate following CS delivery and decreases in rate following US delivery after training. A two-way ANOVA found a significant interaction between training (Early, Late) and stimulus (CS, US) in their effect on peak neural responding (F(1,147) = 87.65, p < 0.0001, n = 153 units from 5 mice). Post-hoc tests found that Early in training, activity after the US was greater than activity after the CS (p < 0.001) and that Late in training, activity after CS was greater than activity after the US (p < 0.001). **D**) Representative change in force of a representative animal as function of training session. **E**) Transitions in CR and UR force from the same sessions as the neural recordings shown in ***B***. **F**) CR force increases as a function of training while the UR amplitude decreases. A two-way ANOVA showed a significant interaction between training (Early, Late) and response type (CR, UR) on peak change in force (F (1,5) = 57.04, p < 0.001, n = 5 mice). Post-hoc tests found that Early in training, UR force was greater than CR force (p < 0.05) and that Late in training, CR force was greater than UR force (p < 0.01). Data represent mean +/− SEM. * p < 0.05, ** p<0.01, *** p < 0.001, **** p < 0.0001.

Several additional features of Forward DA neurons indicated their role in generating movements during the task. For example, regardless of the extent of training, Forward DA neurons were only activated by the CS if the animal made a subsequent CR (**Supplemental Figure 5**). Even well-trained mice occasionally did not show anticipatory movement as measured by force exertion. In such cases, Forward DA neurons were not activated (**Supplemental Figure 5**). Sometimes mice were already engaged in spontaneous movements at the start of the trial, as detected by force sensors, so CS presentation interrupted the spontaneous movements. When this occurs, Forward DA neurons also showed less activity.

With learning the fraction of trials containing a CR increased, and the latency of the force CR became less variable (**Supplemental Figure 5).** The probability of burst activity in DA neurons was correlated with the fraction of trials containing CRs. Training reduced the variance in burst latency following the CR, as the timing of the movements became more consistent. On the other hand, we did not observe any changes in firing rate or bursting probability of Backward DA neurons, probably because training mainly resulted in increased forward movements reflecting anticipatory approach behavior (**Supplemental Figure 6**).

### Force exertion explains how DA firing is modulated by reward probability or size

Phasic DA signaling changes with reward size manipulations^9,11^. Previous studies did not quantify the behavioral changes associated with such manipulations other than licking. We found that manipulations of reward size also altered movements generated by mice (Figure 5A). Increasing reward size resulted in longer duration URs (Figure 5B), and the mean force exerted was greater for both the CR and the UR (Figure 5C). Consistent with prior work, Forward DA neurons increased their firing rates to the CS and to the US with a larger reward (Figure 5D & E). Average firing rates for the Forward DA populations were greater during trials with larger rewards (Figure 5F). On the other hand, Backward DA populations did not respond differently to the changes in reward size (**Supplemental Figure 7**).

**Figure 5.**
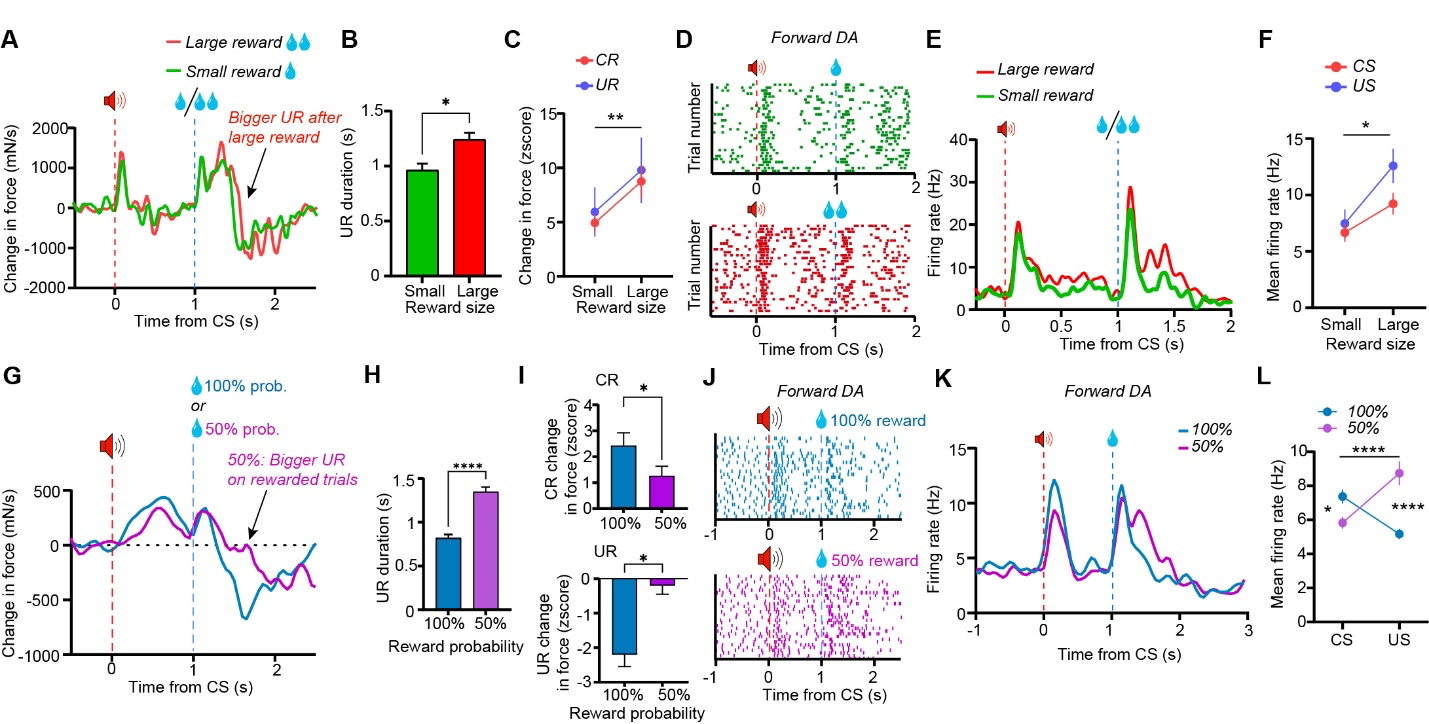
Larger DA responses to greater reward magnitude or uncertainty can be explained by increased force exertion. **A**) Representative changes in force on trials with different reward size. **B**) UR duration is longer on trials with larger reward (paired t-test, p < 0.05, n = 8). **C**) Mice exert greater force during both the CR and UR when larger rewards are delivered (Two-way ANOVA, significant main effect of Reward Size, F(1,6) = 14.63, p < 0.01). **D**) Example raster plots showing the effect of reward size on bursting activity of a Forward DA neuron. **E**) Example mean firing rate of a Forward DA neuron showing effects of reward size on firing activity. **F**) Mean firing rates were greater following larger rewards after both the CS and the US (Two-way ANOVA, significant main effects of Reward size F(1,27) = 5.448, p < 0.05 and Response F(1,27) = 6.953, p < 0.05, n = 23 neurons). **G**) Representative change in force on rewarded trials under different levels of uncertainty. **H**) UR duration is extended on rewarded trials that are less certain (paired t-test, p < 0.0001, n = 6). **I**) Top, mice exert greater force during the CR when rewards are more probable (paired t-test, p < 0.05, n = 6). Bottom, mice show greater changes in force on rewarded trials when reward is less likely (paired t-test, p < 0.05, n = 6). **J**) Left, Raster plots showing firing activity of a representative Forward DA neuron recorded while rewards were delivered with 100% probability and 50% probability. **K**) Extra firing after reward promotes greater force exertion on trials with lower reward probability. **L**) While reward delivery is uncertain, forward DA neurons show lower mean firing after the CS and greater firing after the US. When reward is more reliable, Forward DA neurons fire more after CS, but are not as activated after the US (Two-way ANOVA, significant interaction between trial phase and reward probability, F(1,56) = 55.37, p < 0.0001. Post-hoc tests revealed that firing rate was higher after CS in 100% condition than 50% condition (p < 0.05) but was higher after US in 50% condition than 100% condition, p < 0.0001, n = 30 neurons). Data represent mean +/− SEM. * p < 0.05, ** p < 0.01, **** p < 0.0001.

Prior work also found that phasic DA activity was modulated by reward probability^9^. We manipulated reward probability and compared its effect on force exerted and DA activation (Figure 5G). The duration of the UR was extended when mice received rewards only 50% of the time (Figure 5H). Mice produced lower CR force when reward probability was 50%, but generated larger UR force upon reward delivery (Figure 5I). These patterns were paralleled by similar changes in firing rate of Forward DA neurons (Figure 5J & K). Firing rates after the CS were reduced when the reward probability was 50%. Once mice received the uncertain reward, however, DA neurons showed higher activity. The opposite pattern was observed when the reward was delivered at 100% probability (Figure 5L).

### Dissociating reward prediction error from force tuning

To test whether phasic DA represents reward prediction or force exertion, we manipulated the location of the reward spout while keeping the reward predictability constant. Our head-fixed setup is flexible enough so that the mouse can still access the reward when the spout is moved slightly behind the mouth. Although this change in spout location is only ∼ 2 mm, it required a change in the direction of movement to access the reward (**Supplemental Video 2**). Because the value of the outcome is identical, the RPE hypothesis predicts similar phasic DA regardless of spout location. In contrast, if the DA neurons represent force vectors, DA neurons with different force tuning (e.g. forward and backward) should be differentially recruited when the spout is moved to a new location (Figure 6A).

**Figure 6.**
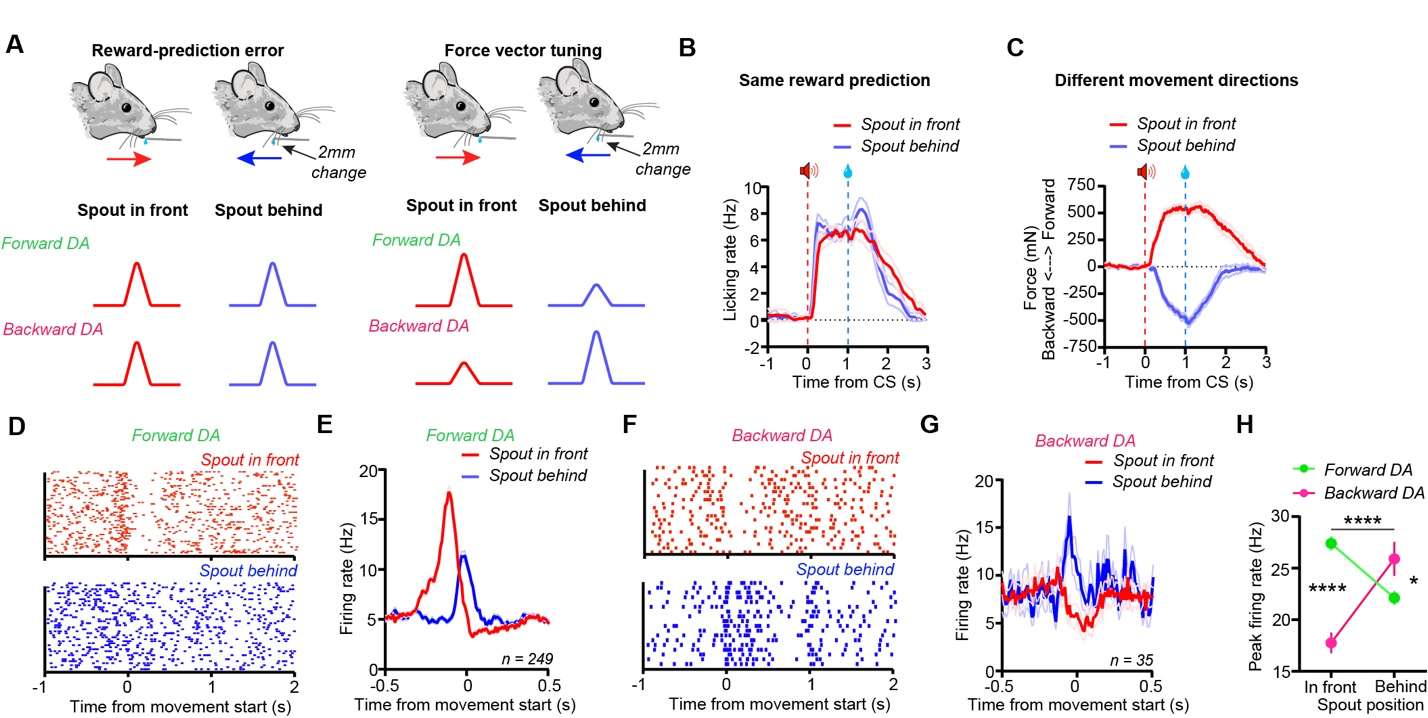
Distinct populations of VTA DA neurons represent force in different directions, despite identical reward prediction. **A)** Force sensors are slightly flexible and allow mice to move small distances forward or backward. The spout was positioned either in front of or slightly under the mice’ chin to encourage distinct movement directions to obtain reward of the same magnitude and value (**Supplemental Video 2**). Diagrams show that firing patterns of Forward and Backward DA neurons are distinguished between the two conditions because of their opponent force tuning. **B**) Anticipatory and consummatory licking rates are identical between ‘spout in front’ and ‘spout behind’ conditions. **C**) Mice make movements in opposed directions between the ‘spout in front’ and ‘spout behind’ conditions. **D**) Example raster plots showing firing of Forward DA neuron aligned to within-task movement under the two spout position conditions (in front vs behind). **E**) Forward DA neurons showed greater responding to movements forward than backward (n = 249). **F**) Example raster plots showing firing of Backward DA neuron aligned to within-task movement under the two spout position conditions (in front vs behind). **G**) Backward DA neurons showed greater firing to movements backward than forward (n = 35). **H**) Opponent activity of Forward and Backward DA neurons depending on reward location (Two-way ANOVA, significant interaction F(1,408) = 83.74, p < 0.0001. Post hoc tests revealed that Forward neurons had firing rates higher than Backward neurons when the spout was in front (p < 0.0001), but Backward neurons had higher responses than Forward neurons when the spout was behind (p < 0.05). Data represent mean +/− SEM. * p < 0.05, **** p < 0.0001.

Anticipatory licking rates (a conventional measure of the CR on this task) were similar for both spout positions (Figure 6B), indicating similar reward prediction regardless of reward location. However, the slight change in spout position resulted in different movements generated by the mice: when the spout was moved backward by only 2 mm, mice generated more backward force and less forward force (Figure 6C). As expected based on force tuning, Forward DA neurons showed higher firing rates when the spout was positioned in front than when it was behind the mouth (Figures 6D **& E**). In contrast, Backward DA neurons showed greater activation when mice were required to exert force in the backward direction (Figures 6F **& G**). These opponent relationships were consistent with how these neurons fired during spontaneous movements (**Supplemental Figure 8**) and further demonstrate that the activity of DA neurons in reward learning tasks are related to online performance rather than learning.

### Phasic DA to aversive stimuli is explained by changes in force direction

If DA activity reflects force exertion, the same relationship between DA and force should be observed regardless of whether the outcome is rewarding or aversive. According to the RPE hypothesis, an unexpected aversive stimulus should produce a negative prediction error, reflected in a decrease or pause in VTA DA activity ^27,28^. To test this possibility, we delivered aversive air puffs on separate sessions (Figure 7A) ^18^. Unexpected air puffs resulted in a movement backwards away from the source of the air puff, followed by rebound movement forward (Figure 7B). This pattern was observed across all mice (Figure 7C). Latency measures confirmed the backward component had a shorter latency than the rebound forward component (Figure 7D). Contrary to the RPE hypothesis, we found that the direction-selective DA neurons reflected the temporal patterns of force changes during an aversive air puff (Figure 7E). The Backward DA population was activated first during the initial backward movement, followed by the Forward DA population that was activated prior to the onset of forward movement (Figure 7F). Forward DA neurons and Backward DA neurons did not differ in their response magnitudes (Figure 7G). Lastly, we also calculated force tuning in these two populations for movements generated after the air puff. Consistent with our prior results (Figure 3), Forward and Backward DA neurons maintained tuning for changes in force in their preferred direction (Figure 7H). Our results agree with past research showing activation of DA neurons in response to aversive stimuli as well as rewarding stimuli, ^3,6,18,29^ but we now show that previous observations could be explained by distinct types of DA neurons signaling force in different directions.

**Figure 7.**
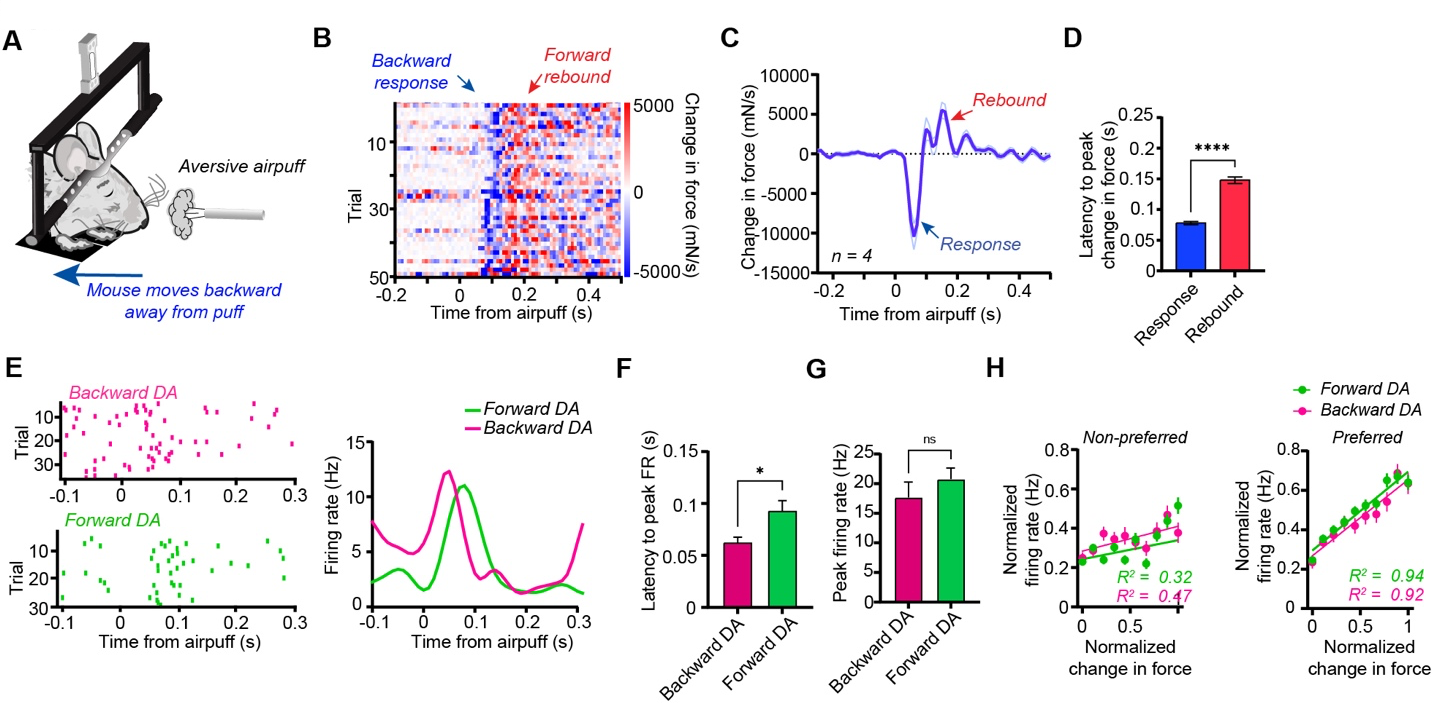
DA activation to aversive air puff produces is explained by bidirectional force changes. **A**) We exposed mice to a mild un-cued air puff delivery to the face in a session separate from Pavlovian conditioning **B**) Mice respond to air puffs with a backward change in force followed by a forward change in force (rebound). **C**) Average change in force trace across mice (n = 4). **D**) Backward responses occurred before the rebound during the air puff trials (paired t-test, p < 0.0001, n = 24 sessions from 4 mice). **E**) *Left,* Rasters for a representative Backward DA neuron and a representative Forward DA neuron showing activation at different latencies. *Right*, average firing rate traces for the neurons whose rasters are shown to the left, note latency differences. **F**) Latency for Backward DA neuron firing was lower than Forward DA (unpaired t-test, p < 0.05, n = 41 Forward DA, n = 35 Backward DA). **G**) Forward DA and Backward DA peak firing rates were not different (unpaired t-test, p > 0.05, n = 41 Forward DA, n = 35 Backward DA). **H**) Forward and Backward DA neuron firing activity was tuned for each population’s preferred force direction during air puff trials (*Left*, Non-preferred, Forward DA, R^2^ = 0.32, p > 0.05, Backward DA, R^2^ = 0.47, p < 0.05; *Right*, Preferred, Forward DA, R^2^ = 0.94, p < 0.0001, Backward DA, R^2^ = 0.92, p < 0.0001). Data represent mean +/− SEM. * p < 0.01, **** p < 0.0001.

We also noticed additional features of DA neuron activity that were inconsistent with RPE interpretations. Some Forward DA neurons exhibited multiple bursts after the reward when mice occasionally generated persistent movements (**Supplemental Figure 9**). We also observed that some Backward neurons exhibit additional ‘hump’-like firing during movements backward under ‘spout behind’ conditions. These observations could only be explained by the pattern of force exertion. However, because they were rare, we did not analyze the results further.

### Reward omission reveals ‘dips’ in DA firing and force

Reduced firing after reward omission is thought to signal a negative RPE (actual reward less than predicted) ^26^. In some well-trained mice, we also omitted the reward after CS presentation (Figure 8A). Forward DA neuron populations showed reduced activity after reward omission (Figure 8B & C). Interestingly, after reward omission, mice abruptly terminated force exertion, and did not generate additional force which is normally observed after reward delivery (Figure 8D). The change in force is therefore negative following reward omission (Figure 8E). This shows that the characteristic ‘dip’ in DA neuron activity after reward omission (Figure 8F) coincides with a pronounced ‘dip’ in force exertion (Figure 8G, **Supplemental Video 3**). Backward DA neurons also showed reductions in firing after reward omission, but they were more modest (**Supplemental Figure 8**), which suggests that mice stop forward force exertion rather than producing a backward movement.

**Figure 8.**
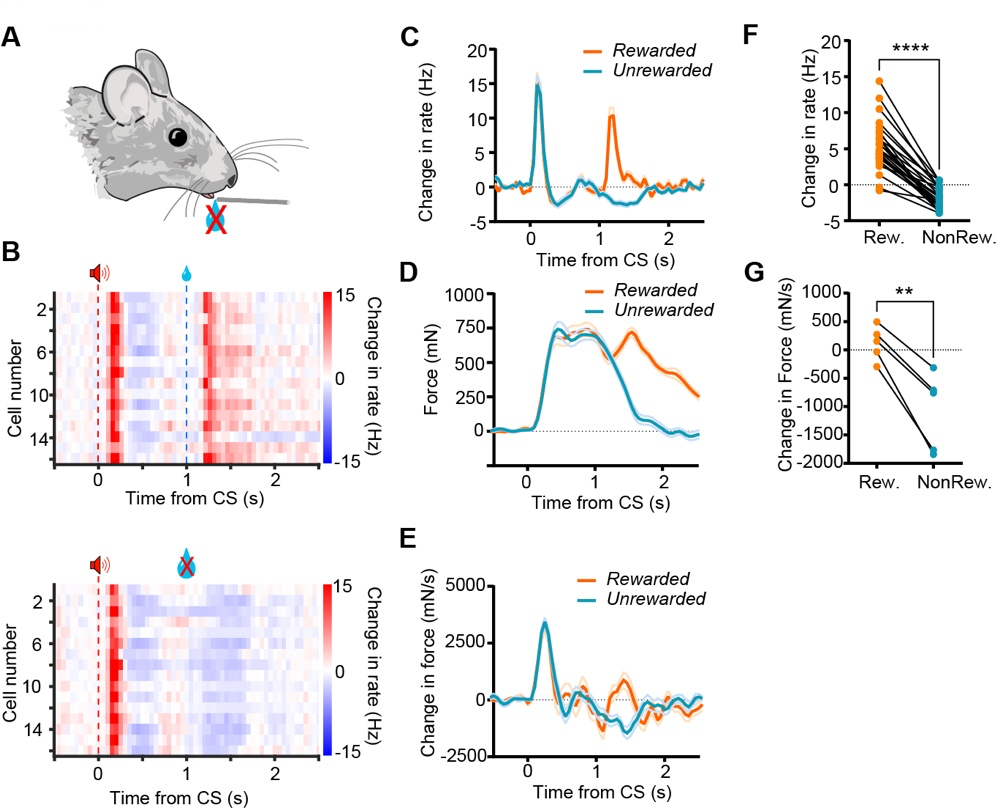
Parallel ‘dips’ in force and DA firing following omission of expected reward. **A)** Schematic of reward omission experiment, whereby expected reward was omitted on 80% of trials. **B**) Simultaneously recorded population of Forward DA neurons during trials with expected reward delivery (*top*) and omission (*bottom*). Note characteristic ‘dip’ in DA neurons during reward omission. **C**) Mean change in firing rate for the population in B during rewarded and unrewarded trials shows a robust dip in activity following omission. **D**) Mice abruptly withhold forward force exertions upon reward omission (**Supplemental Video 3**). **E**) Reward omission results in parallel ‘dips’ in change in force and DA activity. **F**) Reduction in firing rate after reward omission (paired t-test, p < 0.0001, n = 28 neurons from 3 mice). **G**) Omission resulted in a reduction in mean force in the period following reward omission (paired t-test, p < 0.01, n = 5 mice). Data represent mean +/− SEM. ** p < 0.01, **** p < 0.0001.

### Bidirectional optogenetic manipulation altered force exertion without affecting learning

According to the RPE hypothesis, DA serves as a teaching signal to adjust learned associations^2^. We tested whether stimulating DA neurons in place of a sucrose reward after CS presentation would lead to learning of the CS-stimulation association, as reported previously^13^. We trained both VTA *DAT-Cre* with ChR2 expression *(n* = 3) and *WT* mice (n = 5) by delivering a tone (CS) followed by stimulation instead of sucrose reward (Figure 9A). Stimulation did not produce a learned association, as mice did not generate any licking (Figures 9B) or anticipatory forward movement to the CS (Figure 9C). After 400 trials, we then introduced a sucrose reward in addition to stimulation. When compared with control mice, stimulation did not affect the learning rate when the reward was also delivered (Figure 9C). Previous work has shown that activation of DA neurons could produce movement^6,18^. In agreement with such observations, optogenetic stimulation 4 seconds after reward delivery (Figure 9D) also produced forward force (Figure 9E). Moreover, each light pulse resulted in increased force (Figure 9F).

**Figure 9.**
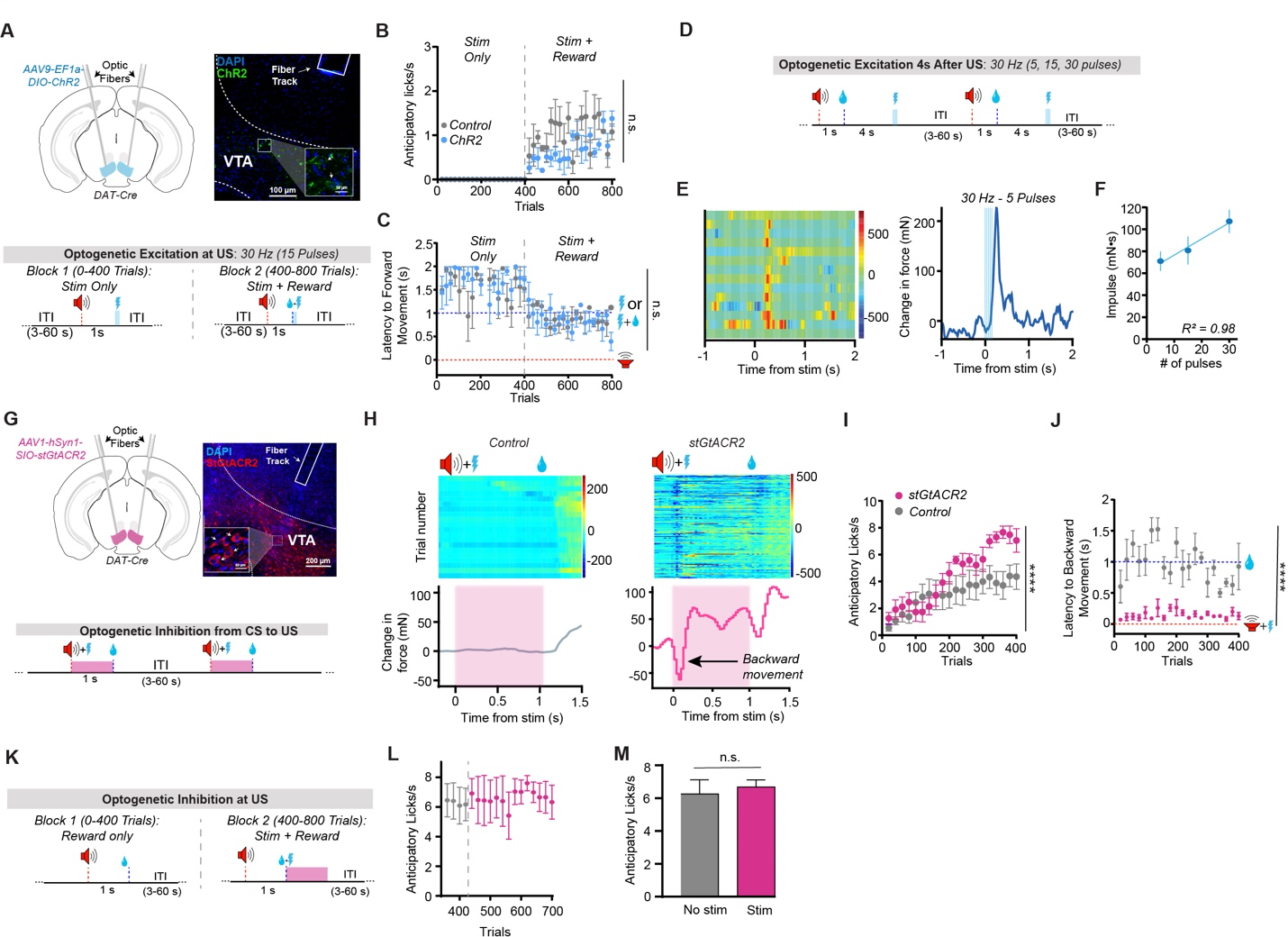
VTA DA is neither necessary nor sufficient for learning the CS-US association. **A**) Cre-dependent ChR2 or DIO-eYFP was injected into the VTA of DAT-Cre mice (N = 3) to excite VTA DA neurons (Top). To test whether VTA DA is sufficient for learning, brief high frequency stimulation (30 Hz – 15 pulses) was used in place of reward (*Bottom*). After 400 trials, a sucrose reward was delivered together with stimulation. **B**) Stimulation did not produce a CS-US association with stimulation alone compared to controls (N = 5), as indicated by anticipatory licking (Two-way ANOVA, significant interaction, F(38,152) = 1.752, p < 0.01, but post-hoc analyses revealed no significant effects between any of the groups (p > 0.05)). **C**) The same was found for latency to move to the CS (Two-way ANOVA, significant main effect of Trial Number, F(38, 152) = 5.80, p <0.0001, no effect of Stimulation F(1,4) = 0.1, no interaction, F(38,152) = 0.9, p > 0.05). **D**) DA neurons were stimulated 4 seconds after reward during a time period when mice are least likely to generate spontaneous movements. **E**) The effect of DA activation on forward force generation. **F**) There is a linear relationship between pulse number and impulse at 30 Hz (R^2^ = 0.98). **G**) We used the Cre-dependent inhibitory opsin stGtACR2 (N = 4) to inhibit VTA DA neurons during learning or DIO-eYFP (N = 6) for controls (Top). Bottom, we inhibited DA activity during the CS-US interval. **H**) Example backward movement resulting from inhibition of DA neurons compared to a control. **I**) Optogenetic inhibition did not prevent learning of the CS-US association, as indicated by anticipatory licking compared to controls. However, it did cause an increase in anticipatory licking, in contradiction with the RPE hypothesis (Two-way ANOVA significant main effects of Stimulation, F(1, 160) = 25.34, p < 0.0001, and of Trial Number, F(19, 160) = 6.775, p < 0.0001, no interaction, F(19,160) = 1.3, p > 0.05). **J**) Optogenetic inhibition significantly reduced latency to backward movement initiation compared to control stimulation (Two-way ANOVA, significant main effect of Stimulation, F(1, 7) = 339.5, p < 0.0001, no main effect of Trial number, F(19, 133) = 0.8, p > 0.05, no interaction F(19,133) = 0.86). **K**) To test whether inhibition causes a negative prediction error, VTA DA neurons were inhibited at the time of reward for 1 second in trained mice. **L**) Inhibition at reward did not significantly impair learning. **M**) There was no change in anticipatory licking following inhibition (paired t-test, p = 0.46). Data represent mean +/− SEM.

To further test whether VTA DA signaling is necessary for learning, we used an inhibitory channelrhodopsin (SIO-*stGtACR2* ^30^, *N* = 4 DAT-cre mice, control: DIO-GFP in N = 6 DAT-Cre mice) to inhibit VTA DA neurons during the CS-US interval (Figure 9G). Inhibition produced backward movement, but did not impair learning (Figure 9H). In fact, mice with DA inhibition licked more after inhibition than controls, contrary to what the RPE hypothesis predicts (Figure 9I). Inhibition also reduced their latency to backward movement compared to controls (Figure 9J). According to the RPE hypothesis, if DA neurons are inhibited immediately after reward delivery, there should be a negative RPE and reduce anticipatory licking on future trials ^31^. Contrary to this prediction, when we inhibited DA neurons at reward delivery to mimic a negative prediction error (Figure 9K), anticipatory licking (CR) was not reduced (Figures 9L **& 9M**). Together our optogenetic results show that phasic DA is neither necessary nor sufficient for stimulus-reward learning.

## Discussion

Using sensitive measures of force generated by head-fixed mice, we demonstrated systematic changes in behavior and the activity of VTA DA neurons during a stimulus-reward task (Figure 1). We found two populations of DA neurons whose activity precedes and predicts force in different directions (Figures 2 **& 3**). This relationship between DA activity and force generated is independent of learning, and can be explained using a simple biophysical model that treats force as proportional to DA concentration in upstream areas (Figure 3, **Supplemental Figure 4**). Although we found the classic pattern of burst firing following reward presentation (US) and the reward-predicting cue (CS), such activity does not signal RPE. Instead, VTA DA neurons regulate performance continuously during reward-guided behavior (Figures 4, **5, 7 & 8, Supplemental Figure 5**) and play a causal role in force generation (Figure 9). By keeping reward prediction constant but changing the direction of movement required by moving the spout, we were able to demonstrate systematic changes in phasic DA activity based on direction of force exertion rather than reward prediction (Figure 6). Force exertion could also explain modulation of DA activity by reward probability or magnitude (Figure 5)^11,32^.

Our results suggest that phasic DA signaling is critical for regulating the direction and magnitude of force exertion continuously, independent of learning. The force tuning of DA neurons was not altered by learning. It was similar during spontaneous movements outside of the task and during movements in response to aversive air puffs. Finally, although it has been proposed that tonic DA activity is related to response vigor ^33^, our results suggest that tonic and phasic components of DA activity represent the same variables, and differ only in magnitude (Figures 2, **3 & 4**).

### Outcome valence

We showed that VTA DA neurons can increase firing to rewarding and aversive stimuli, but these activations are determined by subtle differences in characteristic force profiles (Figures 7, 8, 10), indicating DA activity is independent of outcome valence ^3,29^. A recent study found phasic DA activity in response to aversive events when the probability of receiving a reward was high. In contrast, DA neurons were inhibited when the probability of reward was low ^34^. However, in our study, air puff trials were run separately so the mice only received air puffs without having any probability of receiving a reward. If VTA DA neurons signals RPE, they should be inhibited by the air puff. We found, however, that they were excited and modulated by the direction of movements evoked by the air puff (Figure 7). Importantly, force tuning remains the same despite the change in outcome valence.

### Reward omission

A widely reported observation supporting RPE interpretations of DA is the ‘dip’ in DA firing when mice do not receive an expected reward ^26^. We found that mice abruptly changed the direction of force exertion after reward omission. This negative change in force appears as a ‘dip’ that parallels the dip in DA activity at the expected time of reward delivery (Figure 8).

Recent work also found that distinct populations of VTA DA neurons that project to different sub-regions of the nucleus accumbens (medial and lateral shell) respond differently to aversive stimuli ^35^. However, there was no continuous behavioral measure, and the only behavioral measure during the shock experiments was percent of time spent freezing. It is possible that animals may briefly move in different directions in addition to freezing, as we have found on air puff trials. Thus, VTA DA projections to distinct accumbens regions could be responsible for either forward (approach) or backward (avoidance) movements. In fact, our results suggest that with shock, the activation of opposing force systems may be in conflict and result in no overt movement.

### Limitations of RPE models

Determining how learning impacts performance has long been a major challenge, as it is well established that changes in performance cannot be equated with learning ^4,36^. In reinforcement learning models, the RPE is assumed to be a teaching signal, though in practice learning is often conflated with performance in such models: Reward prediction is assumed to be translated directly into performance, and the RPE that causes ‘learning’ is assumed to change response probability ^37^.

There are two popular interpretations of the DA’s role in reinforcement learning. According to the strong version, DA neurons not only signal RPE’s, but also directly cause learning. According to the weak version, DA activity does not necessarily *cause* learning ^38^. It is only one signal among many in a larger circuit for predictive learning. However, our results are at odds with both accounts. By manipulating reward location while maintaining reward prediction, we tested whether RPE or force generation can better explain DA activity (Figure 6). If phasic DA encodes RPE, reversing the movement direction required to obtain reward should not alter DA signaling, as reward prediction and the value of the reward remain similar as indicated by conventional measures of the conditioned response like anticipatory licking ^3,39^. Although this manipulation changed the location of the reward spout by only ∼2 mm, it significantly changed the direction of force exerted by the animal, as detected by highly sensitive force sensors. As shown in Figure 6, the activity of VTA DA neurons also changed drastically when the reward spout is moved, and this change can be explained by the change in the direction of force exertion. More backward force is associated with increased activity of DA neurons representing backward force, and more forward force increases the activity of DA neurons representing forward force. Neither RPE nor reward prediction can explain this pattern of results.

In addition, the RPE hypothesis fails to predict several other observations, such as multiple successive bursts after reward, and a lack of DA activation to the tone on trials without CRS, even when the mice are well-trained (Figure 2, **Supplemental Figure 5, Supplemental Figure 9**). These observations are compatible with the role of DA in continuous regulation of performance online, rather than updating stimulus-reward associations.

Our optogenetic experiments also indicate that phasic VTA DA is neither necessary nor sufficient for learning (Figure 9). According to the RPE hypothesis, if a burst occurs reliably after a neutral stimulus, the stimulus should generate reward prediction. However, optogenetic stimulation in place of reward did not produce any learning: there was no anticipatory licking, and the rate of learning did not change compared to controls when the stimulation was paired with reward (Figure 9). Optogenetic excitation and inhibition did however produce bidirectional force changes. Force exerted during head-fixation is expected to result in changes in velocity during freely moving behavior ^6,40^.

### Vectorial representation of force

RPEs used in traditional reinforcement learning models are scalar quantities. They do not explain force vector tuning. Tabular reinforcement learning models usually define actions as discrete events, leaving out the spatial and temporal dimensions necessary for generating behavior. With many degrees of freedom, the discretization of the action easily leads to explosive growth of action values and an action space that is too large to explore efficiently. Every possible combination of force direction and magnitude must be learned, and their value stored in memory for the appropriate action to be chosen, which represents an unrealistic computational demand for the nervous system.

Although more recent continuous reinforcement learning approaches attempt to address the limitations of tabular models^41,42^, they still cannot explain our results. For example, using deep-Q learning, Lillicrap et al attempt to generate continuous control signals by mapping states to a probability distribution of torque values. However, millions of learning trials are needed just to generate a simple movement in simulation, and the learned policy is unlikely to work with unpredictable environmental disturbances. Recently, Lee et al argued that a vector RPE model can explain results showing heterogeneous coding of task variables but uniform reward responses^43^. They suggest that DA heterogeneity reflects a high-dimensional state representation with a distributed RPE code. However, their results do not rule out the possibility that the encoding of diverse variables, such as response to cues or movement, can be explained by the same underlying variable of force generation, which they did not measure.

Our findings indicate that DA neurons do not show uniform reward responses. Phasic DA activity at the time of CS, reward, or spontaneous movements can all be explained by the force vector, which we propose is proportional to the amount of DA release in downstream target region that regulate movements in distinct directions, such as the striatum. This is supported by our biophysical model of striatal DA concentration as a function of DA firing rate (Figure 3). By assuming that Forward neurons and Backward neurons project to distinct target regions, we were able to predict the actual force vector using the firing rates of different DA populations.

Further work is needed to test this model experimentally using direct measures of DA in the striatum while recording force changes.

### Limitations in traditional experimental designs and measures

Our results suggest that traditional experimental design and analysis have obscured the key behavioral changes during learning, resulting in misinterpretations of the role of phasic DA in learning and performance (Figure 10). The lack of continuous behavioral measures with high temporal and spatial resolution is a major limitation in previous research on the role of DA in learning. Previous work relied on discrete measures of licking or limb movements ^3,8^. More recent work in head-fixed mice attempted to quantify behavior more thoroughly ^25,44-46^, but still neglected direction-specific torso and head movements, which are critical in understanding the contribution of DA neurons in tasks that are organized around feeding behaviors.

**Figure 10.**
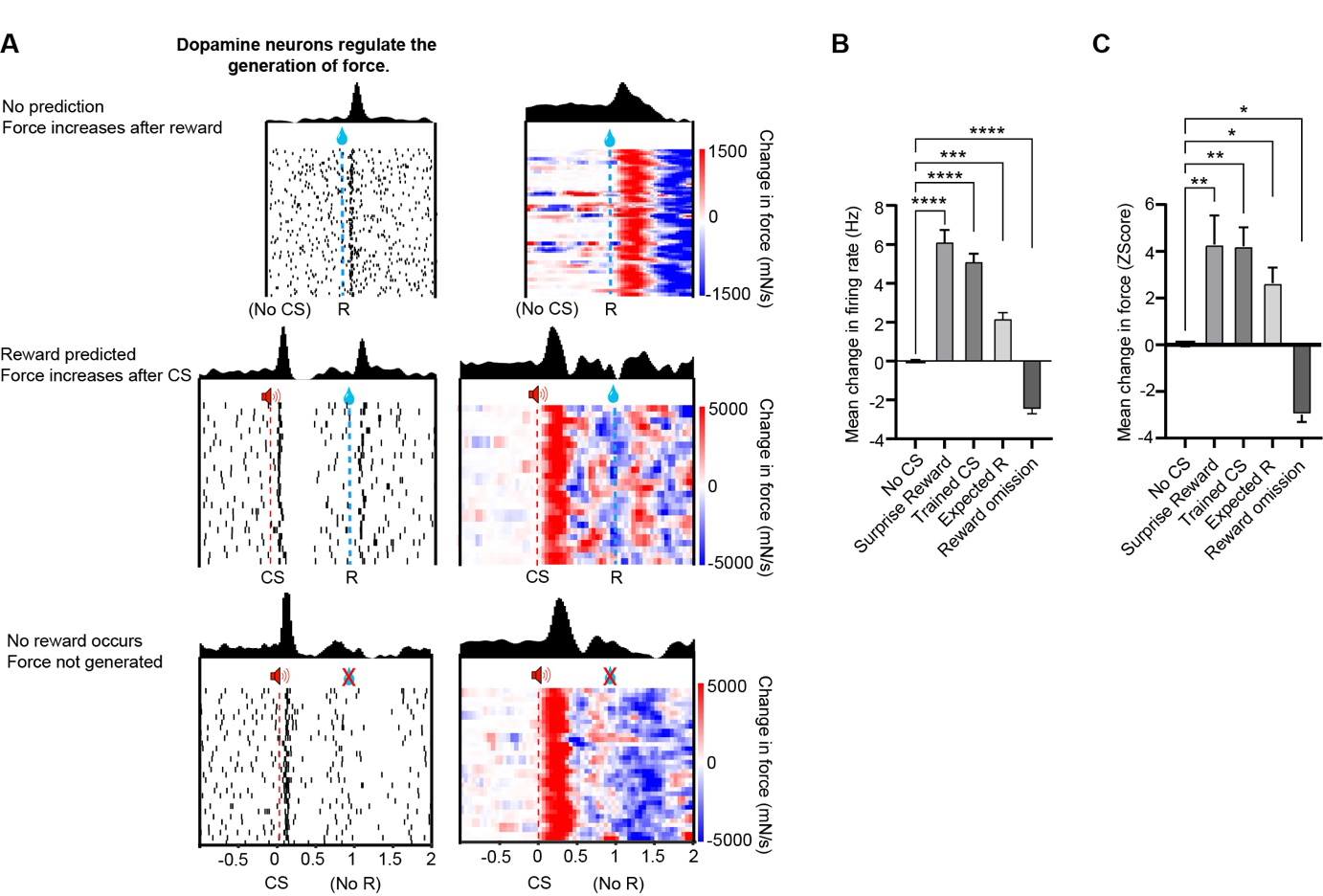
Changes in DA activity explain changes in force exertion rather than reward prediction. **A**) Raster plots of example neurons from mice depicting classically observed DA firing patterns to surprise reward, reward predicted by tone, and omitted reward. On the right are changes in force occurring in parallel with DA responses in the same session. Mice push forward to surprise rewards, to cues predicting reward, and after receiving expected reward. Mice terminate force generation upon reward omission, producing a negative change in force. **B**) Bidirectional changes in firing rate under conditions depicted in A (One-way ANOVA, F(4,131) = 7.971, p < 0.0001; Post hoc tests using No CS condition as the control showed increased firing after Surprise reward, p < 0.0001, after CS in well-trained mice, p < 0.0001, in response to expected reward, p < 0.001, and significant decreases after omission of expected reward, p < 0.0001; n = 84 neurons). **C**) Mean changes in force for the same sessions (One-way ANOVA, F(4,24) = 2.31, p < 0.0001; Post hoc tests using No CS condition as the control showed increases in force after Surprise reward, p < 0.01, after CS in well-trained mice, p < 0.01, in response to expected reward, p < 0.05, and significant negative changes in force after omission of expected reward, p < 0.05; n = 10 mice). Note that patterns of force changes closely parallel changes in DA activity shown in B. Data represent mean +/− SEM. * p < 0.05, ** p < 0.01, *** p < 0.001, **** p < 0.0001.

Earlier work attempted to monitor EMG signals from the arm in restrained monkeys and found no clear relationship between arm muscle and DA activity ^10^. We demonstrate that head-restrained mice attempt to move their heads when attempting to obtain water with the mouth. Measurements of arm muscle activity are inadequate if the key movements are generated by the head and torso. Lacking adequate behavioral measures, previous work could not rule out confounding factors like force exertion and changes in such performance variables during learning.

## Conclusions

By using precise and continuous force measurements, we have shown that VTA DA neurons can carry signals that specify both the magnitude and direction of behavior. Our results suggest that previous results used to support the RPE hypothesis can be explained by a performance account if the force exerted is precisely measured.

During stimulus-reward learning, there was a significant increase in anticipatory force exertion, even though the force tuning of DA neurons remains the same. This observation suggests that the learning mechanism takes place upstream of DA neurons, and the excitatory drive to DA neurons in response to the CS increases with training. In turn, DA signaling may be recruited to modulate the gain of corticostriatal transmission in generating approach behavior. DA concentration in the striatal targets of the mesolimbic DA pathway is expected to reflect force, which is roughly proportional to the integral of phasic DA activity^18^.

Our results are in accord with previous research that has implicated VTA DA in effort, vigor, or incentive salience ^4,5,47-49^. However, these concepts lack detailed vectorial representations needed to explain the spatiotemporal features of continuous behavior. Based on our results, it is expected that DA would produce changes in movement velocity in unrestrained mice ^6,50^. Since force generation is an analog of approach and retreat behavior, and dissociable from licking, our results are also in accord with previous findings that VTA DA contributes more to approach rather than consummatory behaviors ^51,52^. We hypothesize that, by acting as an adaptive gain signal that modulates ventral striatal output, VTA DA regulates the continuous performance of anticipatory approach and avoidance ^53^. This hypothesis can be tested by determining how specific populations of DA neurons exert their effects on target striatal projection neurons.

## Methods

### Subjects

All experimental procedures were approved by the Animal Care and Use Committee at Duke University (Protocol # 162-22-09). 10 DAT-ires-Cre mice, 6 DAT-Cre + Ai32 mice and 8 wild-type male and female mice were used (Jackson Labs, Bar Harbor, ME). DAT::Ai32 mice were generated by crossing Ai32 mice, which express channelrhodopsin (ChR2) in neurons with Cre recombinase, and DAT-Cre mice, which express Cre under the control of the dopamine transporter (DAT) promoter. Mice (2-8 months old) were group housed on a 12:12 light cycle, with experimentation occurring during the light phase. During testing, mice were put on water restriction and maintained at 85-90% of their initial body weights. They received free access to water for approximately two hours following daily experimental sessions.

### Viral Constructs

rAAV9.EF1α.DIO.hChR2(H134R) (Addgene plasmid # 35507 AAV9) and pAAV-hSyn1-SIO-stGtACR2-FusionRed (Addgene viral prep # 105677-AAV1) were used in this study.

### Surgery

Mice were anesthetized with 2.0 - 3.0% isoflurane, and then placed into a stereotactic frame (David Kopf Instruments, Tujunga, CA) and maintained at 1.0-1.5 % isoflurane for surgical procedures. A craniotomy was then drilled above the VTA (AP: 3.1 – 3.4 mm relative to bregma, ML: 0.4 – 0.6 mm relative to bregma, DV: 4.0 – 4.4 mm relative to brain surface). For electrophysiological recordings, drivable electrodes were placed just above the VTA (AP: 3.2-3.4 mm, ML 0.3-0.6 mm, DV, −3.8 mm) and 16-channel recording electrodes were lowered into the VTA (AP: 3.2-3.4 mm, ML 0.5 mm, DV, −4.0 - 4.4 mm).

For optotagging experiments, an optic fiber was attached to the electrode array at an angle (∼15°). For optogenetic experiments, 300 nL of DIO-ChR2 or SIO-StGtaCR2 were bilaterally infused into the VTA of DAT-Cre or WT mice using a microinjector (Nanoject 3000, Drummond Scientific) at a rate of 1 nL per second. The injection pipette was left to sit for 3-5 minutes in order to allow the virus to absorb into the brain tissue and prevent leakage. Custom-made optic fibers (5 - 6 mm length below ferrule, >70% transmittance, 105 μm core diameter) were then implanted at an angle (15°) above the VTA (AP: 3.2 – 3.4 mm, ML: 1.6 mm, DV: 3.8 mm). Fibers and electrodes were secured to the skull using screws and dental acrylic and all mice were fitted with a titanium headbar implant for head fixation. All animals were allowed to recover for two weeks before beginning training on the Pavlovian task.

### Histology

Mice were transcardially perfused with 0.1M phosphate buffered saline (PBS) followed by 4% paraformaldehyde (PFA) in order to confirm viral expression as well as optic fiber and electrode placement. To confirm placement, brains were stored in 4% PFA with 30% sucrose for 72 hrs. Tissue was then post-fixed for 24 hours in 30% sucrose before cryostat sectioning coronally (Leica CM1850) at 60 µm. Fiber and electrode implantation sites were then verified. To confirm eYFP and FusionRed expression in DAT+ cells in the VTA of DAT-ires-Cre and DAT + Ai32 transgenic mice, sections were rinsed in 0.1M PBS for 20 min before being placed in a PBS-based blocking solution. The solution contained 5% goat serum and 0.1% Triton X-100 and was allowed to sit at room temperature for 1 hr. Sections were then incubated with a primary antibody (polyclonal rabbit anti-TH 1:500 dilution, ThermoFisher, catalog no. P21962; polyclonal chicken anti-EGFP, 1:500 dilution, Abcam, catalog no. ab13970) in blocking solution overnight at 4 °C. Sections were then rinsed in PBS for 20 min before being placed in a blocking solution with secondary antibody used to visualize DAT neurons in the VTA (goat anti-rabbit Alexa Fluor 594, 1:1000 dilution, Abcam, catalog no. ab150080; goat anti-chicken Alexa Fluor 488, 1:1000 dilution, Life Technologies, catalog no. A11039) for 1 hr at room temperature. Sections were mounted and immediately coverslipped with Fluoromount G with DAPI medium (Electron Microscopy Sciences; catalog no. 17984-24). Placement was validated using an Axio Imager.V16 upright microscope (Zeiss) and fluorescent images were acquired and stitched using a Z780 inverted microscope (Zeiss).

### Head-fixed behavioral system

The head-fixation device for measuring forces exerted by the animals during behavioral testing and stimulation was described previously ^17,19^. Briefly, the head was clamped via the headbar into the head-fixation frame which contained force sensors (100g load-cells, RB-Phil-203, RobotShop.com). Load cells measure force by linearly translating mechanical deflections into a voltage signal. This voltage signal was then amplified using an INA125P (Texas Instruments) in a circuit configuration that allowed for bidirectional measurement of force. Load cell voltages (1 kHz sampling rate), electrophysiological data, and timestamps for licks, reward, laser were recorded with a Cerebus data acquisition system (Blackrock Microsystems) for offline analysis. A spout connected to a reservoir with a 10 % sucrose solution was positioned at various locations around the mouth. Reward delivery was controlled by opening a solenoid valve (161T010, NResearch, NJ) attached to the tubing connected to the spout. A capacitance-touch sensor (MPR121, AdaFruit.com) attached to the spout was used to detect licks.

### Pavlovian stimulus-reward task

Mice were first allowed to habituate to the head-fixed condition for 5 minutes for approximately 1-2 days. Once mice were habituated to head-fixation, mice were trained on approximately 70-100 trials a day to avoid satiety effects. Sessions occurring before the first session where animals generated CRs (both licking and movement) on more than 75% of trials were classified as ‘Early’. Otherwise, sessions were classified as ‘Late’. A spout that delivers 10 % sucrose was positioned in front of the mouth. White noise was continually present in the background. At the beginning of each trial, a 3 KHz tone that lasted 200 ms was presented followed by delivery of 5-30 ul of sucrose 800 ms after the end of the tone. The delay between onset of the tone and reward delivery was 1s. There was a random intertrial interval that varied to prevent any anticipation of the onset of the tone (4-60 s).

Air puff delivery was delivered using an EFD 1500 XL pneumatic fluid dispenser. The puff lasted 20 ms and the output tube was aimed at the face. Air puff trials were conducted as distinct sessions outside of the cue-reward sessions. There was a 5-minute break between the end of a reward session and the beginning of an air puff session.

For Spout Behind sessions, the spout tip was moved slightly underneath (2mm change in position, **Supplemental Video 2**) the chin of the mice so that they had to move backwards to obtain water reward. For reward probability manipulations, mice were run in blocks of 50 trials that alternated between 100% and 50% reward probability, starting with 100% reward probability. For reward magnitude manipulations, sessions contained either small rewards (20 msreward duration, 5µl) or large rewards (60 ms reward duration, 15 µl).

### Wireless in Vivo Electrophysiology

Drivable electrodes were single-drive movable micro-bundles of tungsten electrodes (1 x 16; 23 μm diameter) placed within a guide cannula (Innovative Neurophysiology, Inc.). Electrophysiological data were recorded using a miniaturized wireless head stage (Triangle Biosystems) that was interfaced with a Blackrock Cerebus data acquisition system (Blackrock Microsystems). A digital bandpass filter was applied to the electrophysiological data (250Hz – 5kHz) and spike timestamps and waveforms were recorded at 30kHz. Filtered data were sorted using Offline Sorter (Plexon). A 3:1 signal-to-noise ratio, and an 800 μs or greater refractory period were required for the neural data to be used for analysis. Single units were selected based on a principal component analysis of waveforms using 2 principal components. To obtain new neurons with driveable electrodes, the electrodes were lowered by 100 μm after each behavioral session. All peri-event raster plots were generated using NeuroExplorer (Nex Technologies).

### Optogenetic identification of VTA DA neurons

Several of our functional groups had wide waveforms and low firing rates, indicating they were DA neurons (**Supplemental Figure 3**). Prior work has shown however that DA neurons can produce a low rate of misleading waveform and firing rate properties^54,55^. To overcome these limitations, we used optogenetic tagging to confirm DA neuron identity. We attached fiber optic implants to our drivable electrodes^17^. Only fibers with ≥ 70% light transmittance measured through the optic fiber tip (PM120VA, ThorLabs) were used. Light (5-8 mW, 5 ms pulse width, 10-20 Hz for 1 second for ChR2, 500 ms pulse, 5 mW for inhibition) was delivered via laser (470 nm DPSS laser, Shanghai Laser & Optics) at the end of each behavioral session. Neurons were classified as tagged if each pulse of light produced a spike occurring with a latency of ≤ 6ms and on ≥70% of trials (**Supplemental Figure 3**). A total of 1285 neurons were recorded, of which 159 were tagged using optogenetics.

### Classification of VTA DA neurons

Recent work found that VTA DA neurons can be identified based on their responses to forward and backward movements ^18^. Because responses during Pavlovian conditioning change with learning, we employed a new functional classification scheme to classify VTA DA neurons by comparing the neural activity of all recorded neurons during spontaneous forward and backward movement events, which were a consistent feature of mouse behavior independent of learning (Figure 1, Figure 2, **Supplemental Figure 1**, & **Supplemental Figure 3**). This approach allowed us to identify movement selectivity that occurs independently of Pavlovian conditioning.

For each unit, firing rates were estimated in 2 second windows around the start of spontaneously generated forward and backward movements using 10 ms bins and baseline subtracted by the average of all pre-event baseline firing rates (first 50 bins). We used agglomerative clustering which resulted in 6 different response profiles. All preprocessing was done in Matlab and clustering was performed with the function *clusterdata* using Euclidean distance and ward linkage parameters. After clustering was complete, cells were manually checked for the consistency of the responses. Some neurons were found to be clustered incorrectly and were then manually changed to the appropriate cluster.

### Optogenetic Parameters

Optogenetic stimulation sessions were identical to the Pavlovian conditioning tasks with the electrophysiological recordings, as described above. Pulses of light (Excitation: 5-8mW measured at the tip of the optic fiber connected to the optic implant, 5 ms pulse width, 30 Hz 5-30 pulses; Inhibition: 5 mW, 1 sec pulse width) were delivered via a laser (470nm DPSS laser, Shanghai Laser & Optics) and controlled through an Arduino.

### Force conversions

Force conversion of load cell signals was described previously ^17,19^. Briefly, we calibrated the load cell circuits using a conversion factor (expressed in Newtons per Volt) determined by the linear relationship between the voltage changes resulting from known masses placed on the sensor. Force was determined by multiplying voltage signal by the conversion factor to obtain a value in Newtons. Impulse was calculated using the Matlab function *trapz* to integrate the total area under the force curve over time from movement onset to movement termination^18^. The change in force was the first derivative of the force signal, calculated using the *diff* function in Matlab. The resulting values were divided by the bin size (1 ms) and smoothed by convolving the signal with a gaussian filter with a standard deviation of 12 bins.

### Detection of movement initiation

Forward movements were defined during force exertions that exceeded a threshold of 0.5 standard deviation of force greater than 0, lasted longer than 100 ms and was also separated by at least 100 ms from another movement. Backward movements were defined as events lower than 1.5 standard deviations of all force values below 0. These events had to last longer than 100 ms, had to be separated by at least 500 ms. Backward movements had to occur independently of forward movements and so could not coincide with the end of any forward movement, as sometimes mice moved backward immediately after moving forward.

### Analysis of force tuning

Change in force during spontaneous movements (−500 ms to + 500 ms around movement initiation) was downsampled into 50 ms bins. Firing rate for each neuron was also binned into the same bins, creating two vectors of equal length. A cross-correlation was computed between the firing activity vectors and the change in force for the neuron’s preferred direction of force exertion to determine the time shift between the two signals. Neural activity was then shifted according to the lag between its activity and change in force in the preferred direction. We next sorted the change in force vector according to magnitude and excluded outliers (1^st^ and 99^th^ percentile) and identified 10 evenly spaced monotonically increasing force bins that evenly spanned the range force changes. Neural activity that was obtained concurrently with forces belonging to each bin was averaged. This same procedure was used when analyzing the relationships between single unit activity with force in the non-preferred direction, but using the same lag as was determined from cross correlation with force in the preferred direction. Tuning was normalized to account for different ranges of force and firing rates produced by different animals and cells and in different sessions, respectively.

### Computational model of DA concentration

We assume that Forward and Backward VTA DA neurons have different target cells that contribute to force generation in a preferred direction. Therefore, to approximate the resulting force, we plot the difference between predicted DA concentrations resulting from activity in these two DA populations. A previous model describes DA concentration dynamics in the striatum as follows:

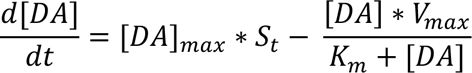

where *[DA]* is the total DA concentration at time t (in µM), *[DA]_max_* is the instantaneous increase in DA concentration every time a neuron spikes (in µM), *S_t_* is the average number of total spikes at time *t*, *V_max_* is the maximal reuptake rate of DA in the striatum, and *K_m_* is the Michaelis-Menten parameter that is the reciprocal of the affinity of the DA transporter. Value constraints for all parameters were determined using prior experimental and modeling work ^21–24^. We neglect the contribution of diffusion on fading DA concentrations, and only consider reuptake in our model. Nothing was tuned to fit the data and no filters was applied. All curves were set at the same average starting baseline and the model curves are scaled so that they have the same maximum as the experimental data.

### Quantification and Statistical Analyses

All analyses were performed with Matlab, NeuroExplorer, and Graphpad Prism. Statistical analyses were performed in Graphpad Prism. A power analysis was not conducted a priori to determine sample size.

## Acknowledgments

We would like to thank Fengxia Allen, Joseph Barter, Guozhong Yu, and Jinyong Zhang for technical assistance. This paper is dedicated to the memory of our co-author Ryan Hughes (1987-2021).

## Author contributions

KIB, RNH, and HHY designed the experiments. RNH and KIB performed surgeries, *in vivo* electrophysiological and optogenetic experiments, and histological analysis. KIB, RNH, IPF, and QJ performed data analysis. MH and BG built the model. KIB, RNH, and HHY wrote the manuscript. All authors have read and approved the manuscript.

