## Supplementary data for "Force tuning explains changes in phasic dopamine signaling during stimulus-reward learning"

### Supplemental Figure 1

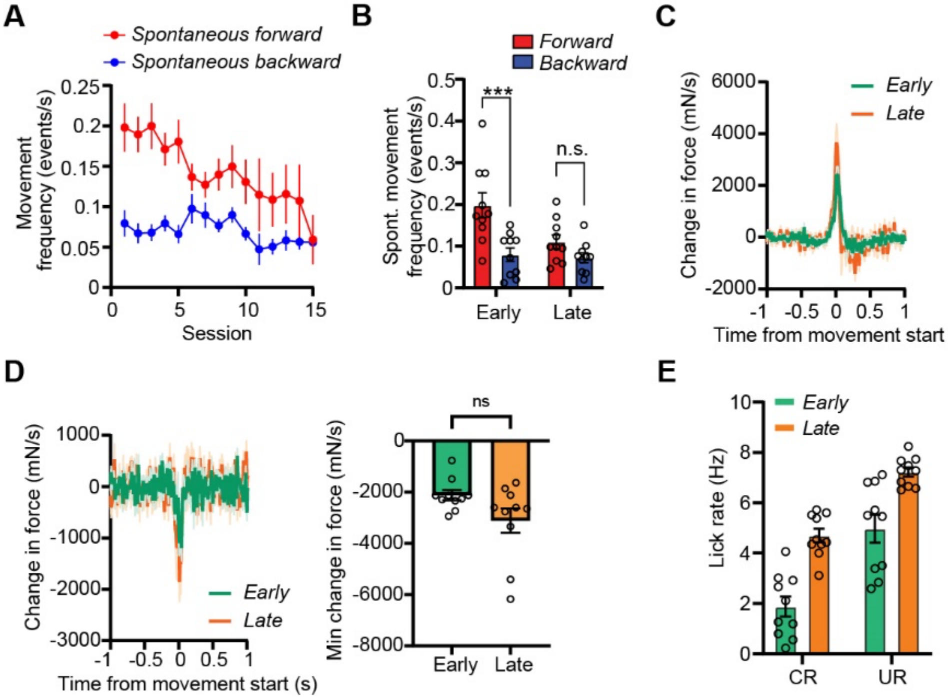


**Supplemental Figure 1. Spontaneous force exertion changes in frequency but not magnitude during learning**

**A**) Mice made less spontaneous forward movements as training progressed. **B)** A two-way repeated measures ANOVA revealed a significant interaction between training stage and movement direction on the rate of spontaneous movement generation ($F(1,9) = 10.05$, $p < 0.05$). Post-hoc tests indicate that mice produce spontaneous forward movements more frequently than backward movements early ($p < 0.001$) but not late ($p>0.05$) in training. **C**) The magnitude of spontaneous forward movements remained consistent across learning (paired t-test, $p > 0.05$). **D**) We did not observe any change in backward force magnitude across training sessions (paired t-test, $p > 0.05$). **E**) Two way ANOVA showed significant effects of task component (CR vs UR, $F(1,9) = 273.2$, $p < 0.0001$) and training stage (Early vs Late, $F(1,9) = 64.3$, $p < 0.001$) on licking rate. No interaction was detected ($F(1,9) = 2$, $p > 0.05$). Data points represent mean +/- SEM. *** $p < 0.001$.

Supplemental Figure 2

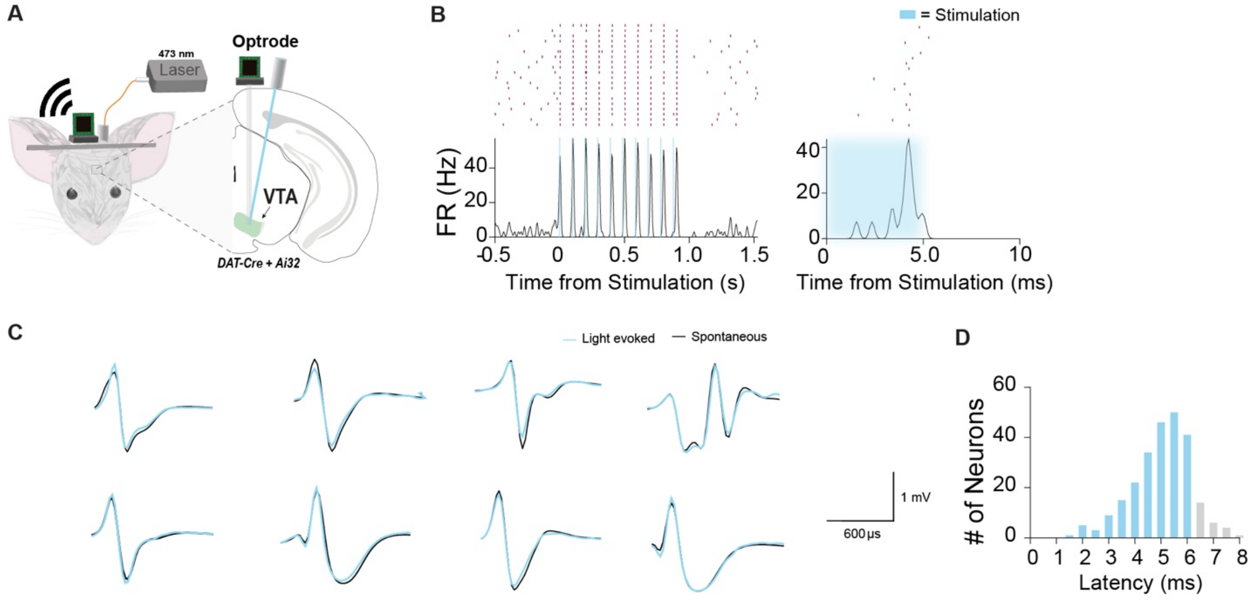


**Supplemental Figure 2. Optogenetic tagging of VTA DA neurons**
**A)** Schematic of optrode placed into the VTA for optogenetic identification of DA neurons during a Pavlovian conditioning task. **B)** Peri-event raster plot of a representative optically tagged DA neuron. **C)** Eight representative optically-tagged DA neuron waveforms that show the same spontaneous and light-evoked waveforms. **D)** Neurons were classified as DA if they had a latency of ≤ 6ms (*n* = 160).

**Supplemental Figure 3**

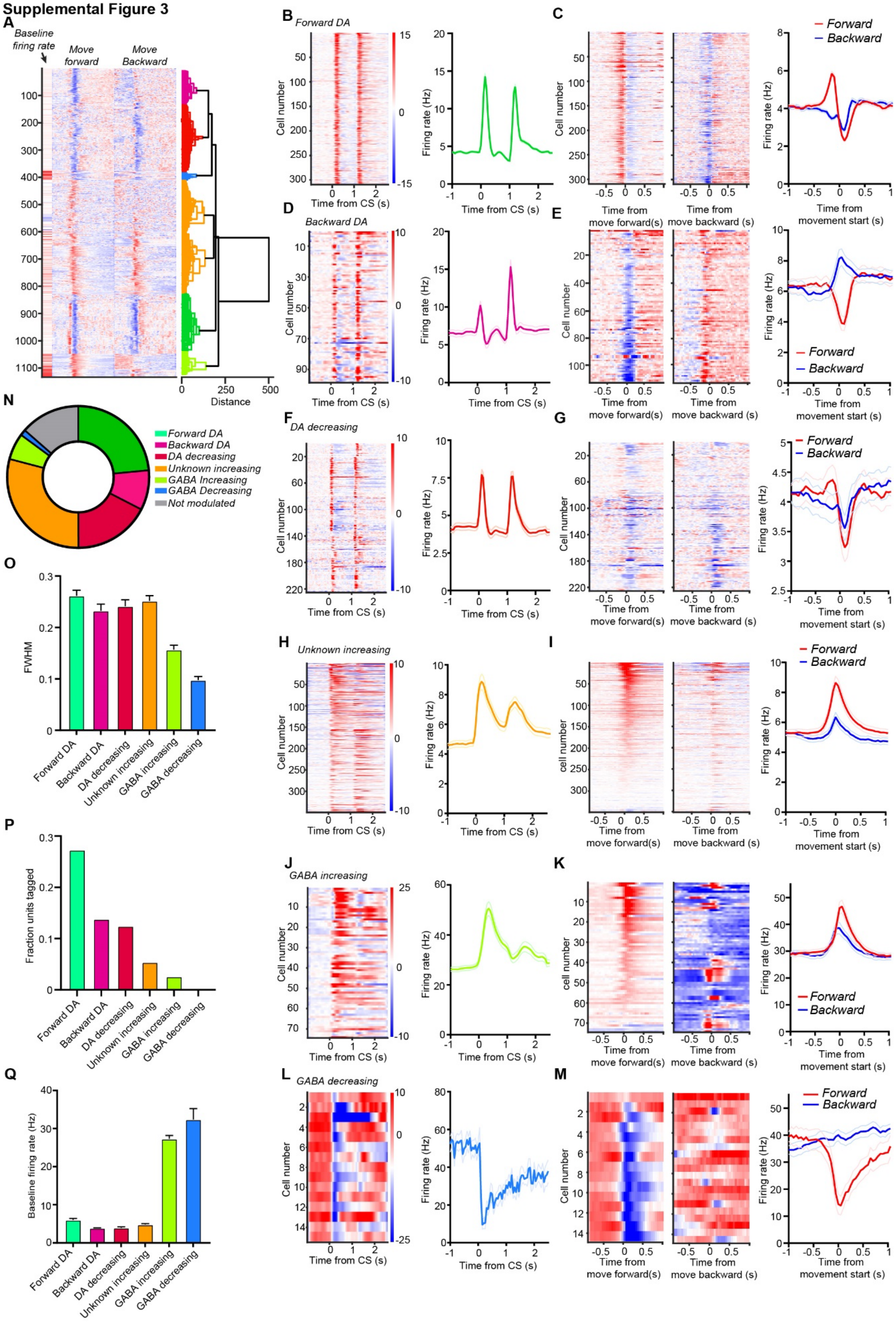

**Supplemental Figure 3. Distinct populations of VTA neurons were found using hierarchical clustering**

**A)** The responses of each recorded neuron to spontaneous (outside of the CS-US interval) forward movements and spontaneous backward movements were normalized using baseline subtraction followed by division by the max to scale firing between 0 and 1. These matrices were then concatenated together along with their baseline firing rates. The resulting matrices were clustered using hierarchical clustering. Three populations were identified as DA based on their proportions of tagged DA neurons, baseline firing rate, and functional responses to spontaneous movements. **B)** Forward DA neurons had bursting activity to the CS and US typical of DA neurons. **C)** Forward DA neurons showed higher responses to forward movements. **D)** Backward DA neurons showed bursting type responses to the task. **E)** Backward DA neurons showed higher responses to movements backward. **F)** The third population of putative DA neurons showed burst -type responding to the task and had relatively high rates of tagging. **G)** This population showed only reductions in firing during movements forward. **H)** An unknown population of neurons had wider, non-bursting responses to the task, but showed lower baseline firing rates and did not show high rates of tagging. **I)** This unknown population increasing in rate to both forward and backward movements. **J)** Of the putative GABA populations, one of them showed increasing firing during the task. **K)** The increasing GABA population showed increasing firing rates to both forward and backward movements. **L)** A second type of GABA population showed reductions in firing during the task. **M**) The decreasing GABA population showed reduced firing during spontaneous forward movements and was not modulated by spontaneous backward movements. **N)** About 50% of recorded neurons belonged to putative DA populations (Forward, Backward, and Decreasing). The remaining neurons were either putative GABA or a population that had DA-like waveforms but rarely tagged. **O)** 4 groups of neurons had wider waveform widths. **P)** Forward, Backward, and Decreasing DA neurons had tagging probabilities of ~12% or higher. The remaining populations showed minimal tagging probabilities. **Q**) Baseline firing rates of different neuronal populations. Putative GABAergic neurons had high baseline firing rates. Data points represent mean +/- SEM.

### Supplemental Figure 4

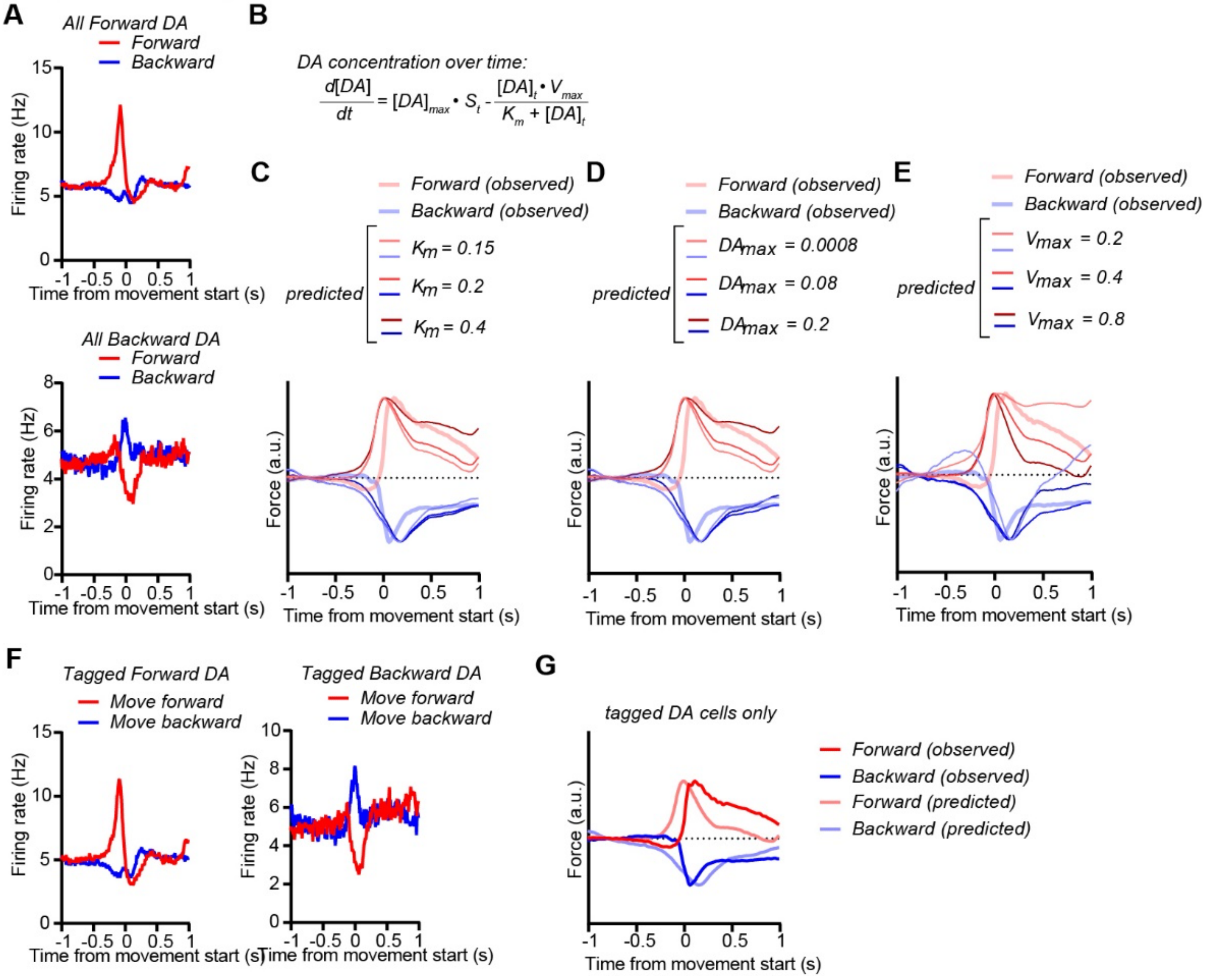


**Supplemental Figure 4. Effects of varying DA concentration parameters on predicted force**

**A**) Mean firing rates for all Forward DA neurons (*Top*) and Backward DA neurons (*Bottom*) in response to movements forward and movements backward. **B**) Equation modeling change in DA concentration during DA spiking activity. **C**) Effects of varying the value of $K_m$, the reciprocal of affinity of the DA transporter for DA, on the model's prediction of exerted force. Lower affinities for DA result in slower decay of force. **D**) Effects of varying the value of $DA_{max}$, the increase in DA concentration for each spike, on the model's prediction of exerted force. More DA release results in slower decay of force. **E**) Effects of varying the value of $V_{max}$, the maximal reuptake rate of the DA transporter, on the model's prediction of exerted force. Less reuptake rates result in slower decay of force. **F**) Mean firing rates for all tagged Forward DA neurons (*Left*) and all tagged Backward DA neurons (*Right*) for forward and backward movements. **G**) The model could predict force exertion using activity from only tagged Forward and Backward DA neurons.

**Supplemental Figure 5**

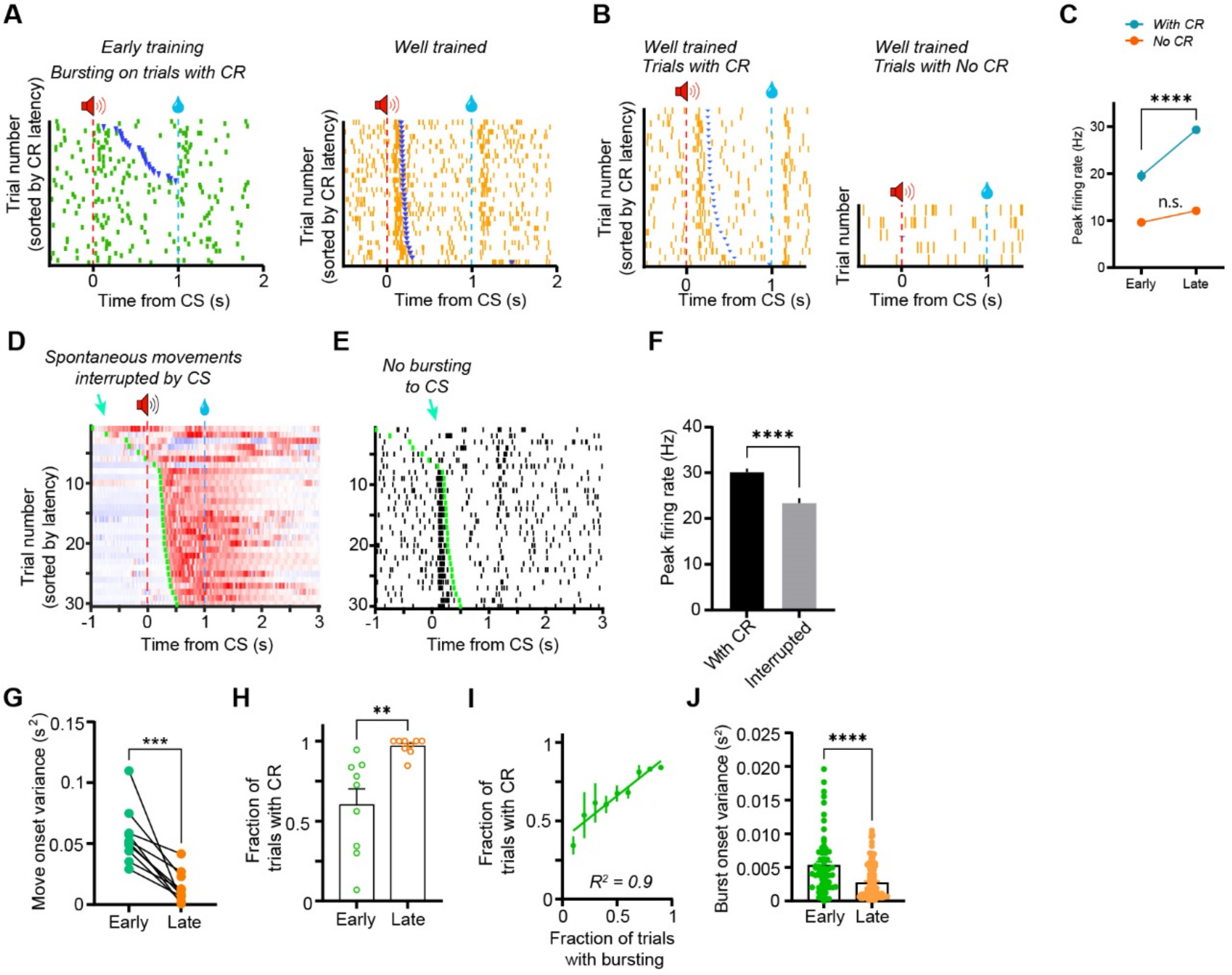


**Supplemental Figure 5. Increased phasic Forward DA activity can be explained by changes in CR timing**

**A)** Example Forward DA neuron rasters sorted by latency to CR onset time (blue dots). **B**) Example Forward DA neuron recorded from well-trained mouse. Despite training, trials lacking a CR did not show bursting in response to CS presentation. **C**) Trials without CR do not alter firing rate changes over learning (Two-way repeated measures ANOVA shows a significant interaction between presence of CR and extent of training on peak firing $F(1,268) = 11.31$, $p < 0.001$; Post-hoc tests showed that Forward DA neurons increase firing only to trials containing a CR with training: with CR, $p < 0.0001$, without CR, $p > 0.05$). **D**) Because trial delivery was random, animals sometimes were moving spontaneously at the time of CS presentation. **E)** Example Forward DA neuron showing reduced responding to the tone it interrupted ongoing movement. F**)** Forward DA responding to CS was lower if animals were occupied with spontaneous movements during CS presentation (unpaired t-test, $p < 0.0001$). **G**) Learning resulted in a significant reduction in movement onset time variance (paired t-test, $p< 0.001$). **H**) The probability of observing a CR increased with training (paired t-test, $p < 0.01$). **I**) The fraction of trials containing bursting activity in Forward DA neurons predicted the rate of CR generation ($R^2 = 0.9$, $p < 0.001$). **J**) Latency for Forward DA neurons to burst after the CS became smaller with training (Unpaired t-test, $p < 0.0001$). Data points represent mean +/- SEM. ** $p < 0.01$, *** $p < 0.001$, **** $p < 0.0001$.

**Supplemental Figure 6**

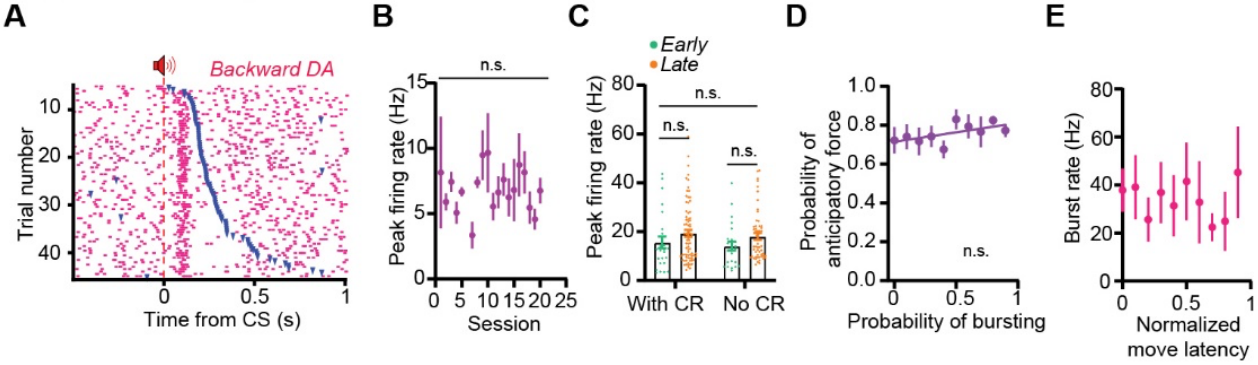


**Supplemental Figure 6. Backward DA neurons are not altered with learning**

**A**) Example raster plot of a Backward DA neuron that bursts to the CS. Bursts do not track the onset of the CR (blue tick marks). Note reduction of firing with forward movement onset. **B**) There was no change in Backward DA peak firing rate in response to the CS over training (One way ANOVA, $p > 0.05$). **C**) The presence of a CR did not increase the peak firing rate of Backward DA neurons, even in later stages of learning (two-way ANOVA, no significant effects of training ($p > 0.05$), CR status ($p > 0.05$), and no interaction ($p > 0.05$). **D**) Bursting activity of Backward DA neurons did not correlate with the likelihood of observing anticipatory force ($R^2 = 0.36$, $p > 0.05$). **E**) Movement latency was not correlated with the firing rates in bursting events of Backward DA neurons ($R^2 = 0.01$, $p > 0.05$). Data points represent mean +/- SEM.

### Supplemental Figure 7

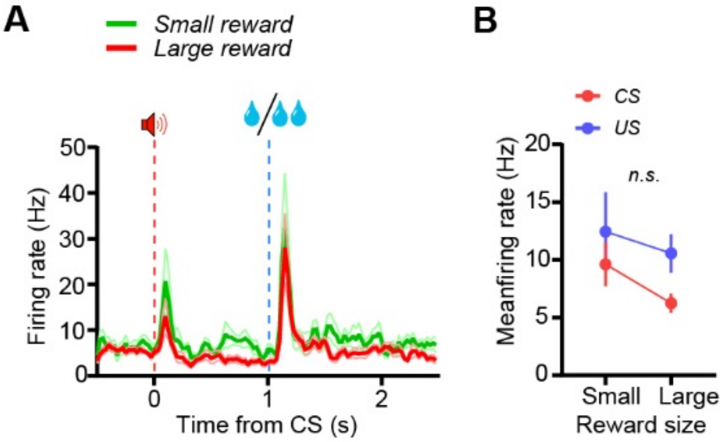


**Supplemental Figure 7. No effects of reward size on Backward DA neurons.**

**A**) Average firing rate of Backward DA neuron population in response to sessions with small or large rewards. **B**) There was no effect of Reward Size on Backward DA firing (Two-way ANOVA, $F(1,15) = 0.31$, $p > 0.05$, $n = 17$ neurons). Data points represent mean +/- SEM.

### Supplemental Figure 8

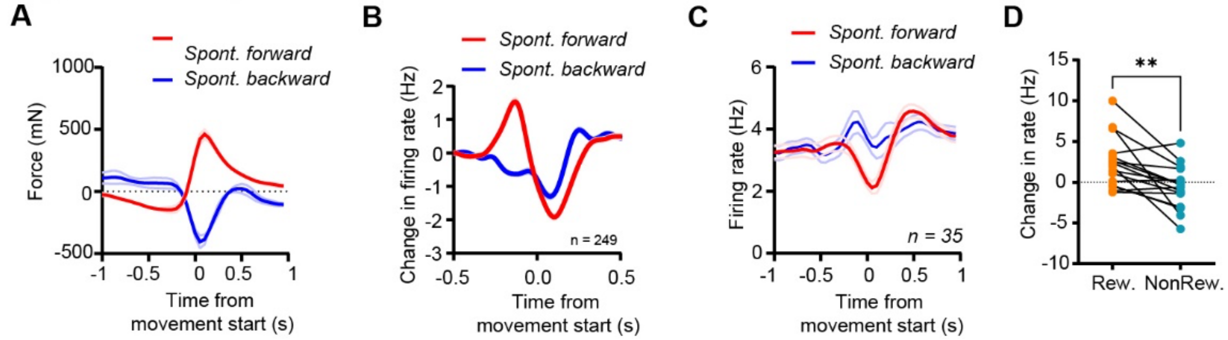


**Supplemental Figure 8. Responses of Forward & Backward DA neurons from Figure 6 during spontaneous movements**

**A)** Mean spontaneous forward and backward change in force for sessions with forward and backward spout positions. **B)** Forward DA neurons had higher responses to spontaneous forward movements **C)** Backward DA neurons showed higher responses to spontaneous backward movements. **D**) Backward DA neurons reduce their firing rate following reward omission (paired t-test, $p < 0.01$). Data points represent mean +/- SEM. ** $p < 0.01$.

### Supplemental Figure 9

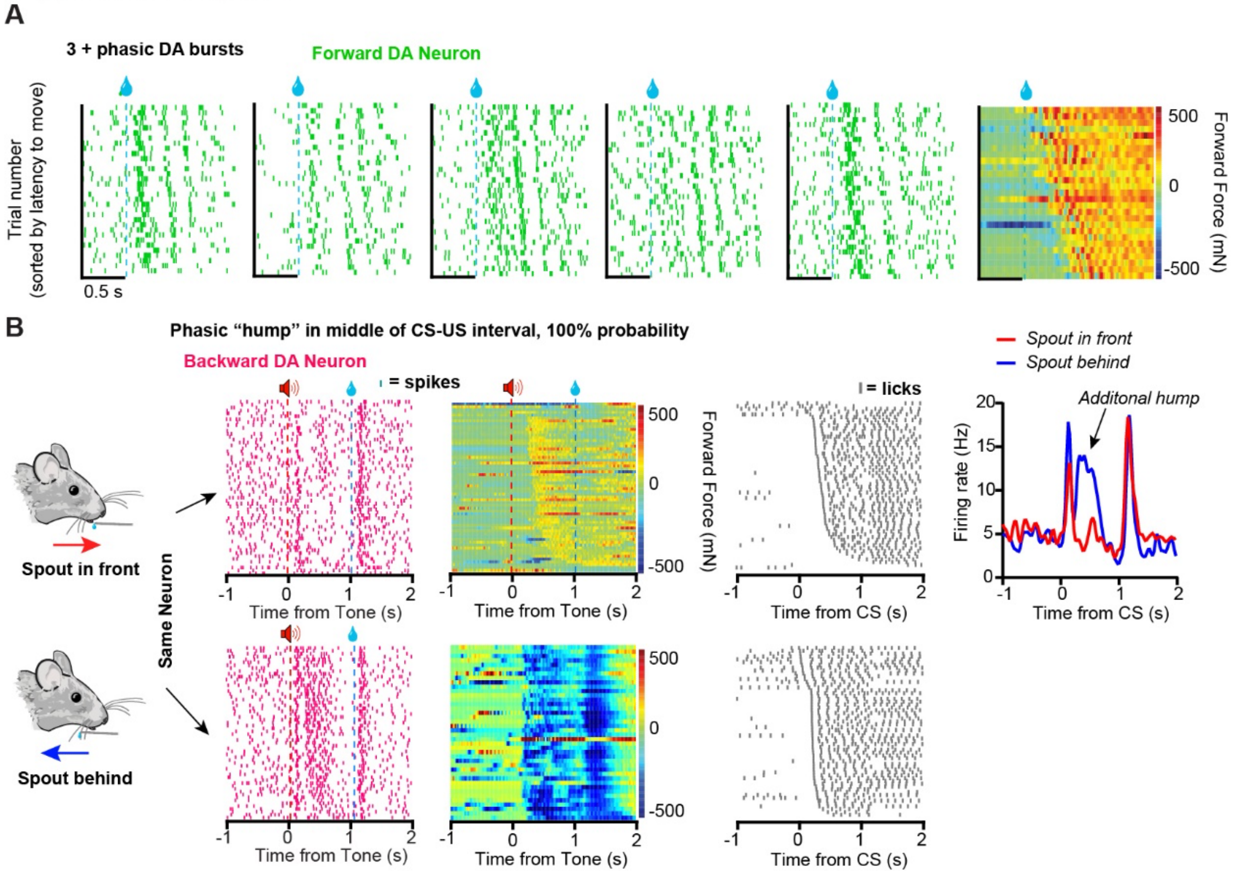


**Supplemental Figure 9. Examples of firing patterns of Forward and Backward DA neurons that are inconsistent with the RPE hypothesis**

**A)** Five Forward DA neurons show up to 4 bursts after the US when forward force was sustained for longer than 1 second. The force sensor indicated corresponding oscillations in force exerted.

**B**) Backward DA neuron displaying a ‘hump’ during backward movement, but not during forward movement. This occurred despite comparable reward prediction, as indicated by anticipatory licking (*middle*). *Right*, average firing rate traces of the example neuron during Spout in front and spout behind conditions.

**Supplemental Video 1. Head-fixed mice generate force during Pavlovian conditioning**

**Supplemental Video 2. Opposing exertion of force with small changes in spout location**

**Supplemental Video 3. Changes in force exertion after omission of expected reward**
